# Visualizing the transcription and replication of influenza A viral RNAs in cells by multiple direct RNA padlock probing and *in-situ* sequencing (mudRapp-seq)

**DOI:** 10.1101/2025.03.26.645430

**Authors:** Shazeb Ahmad, Jianhui Li, Joél Schaust, Anne-Sophie Gribling-Burrer, Nina Geiger, Sabine C. Fischer, Uddhav B. Ambi, Simone Backes, Markus J. Ankenbrand, Redmond P. Smyth

## Abstract

Influenza A viruses (IAV) contain eight negative-sense single-stranded viral RNA (vRNA) molecules, which are transcribed into mRNA and replicated via complementary RNA (cRNA). These processes are tightly regulated, but the precise molecular mechanisms governing the switch from transcription to replication remain elusive. Here, we introduce multiple direct-RNA assisted padlock probing in combination with *in situ* sequencing (mudRapp-seq) to visualize the transcription and replication of all eight IAV vRNA and mRNA molecules at the single-cell level. We demonstrate that direct RNA padlock probing is three times more efficient than conventional probes that target cDNA. Individual probes showed variations in efficiency, partly due to the RNA structure of the target, which was mitigated by employing multiple padlock probes per target. Applying mudRapp-seq to an infection time course, we observed early mRNA expression, followed by vRNA accumulation approximately 3 hours later. Individual viral segments exhibited differential expression, particularly in the mRNA population. Both bulk and single-cell analyses revealed a correlation between the expression of ‘M’ mRNA and the onset of the transcription-to-replication switch. Our findings demonstrate that mudRapp-seq offers significant potential for elucidating viral replication mechanisms and may be applicable to studying other RNA viruses and cellular RNA processes.

## Introduction

Influenza A viruses (IAVs) are important human pathogens responsible for seasonal epidemics and occasional pandemics. They are enveloped viruses with segmented genomes comprised of eight viral RNAs (vRNAs), encoding three RNA polymerase proteins (PB2, PB1, and PA), a nucleoprotein (NP), the surface proteins hemagglutinin (HA) and neuraminidase (NA), matrix (M) and non-structural proteins (NS) (1). Each vRNA associates with the heterotrimeric polymerase complex (PB2, PB1, and PA) and multiple copies of NP to form viral ribonucleoproteins (vRNPs). Unusually for an RNA virus, vRNPs replicate in the nucleus and are only later exported into the cytoplasm for incorporation into nascent particles at the plasma membrane (2). As the vRNAs are in negative-sense, they must be transcribed into positive-sense messenger RNA (mRNA) for translation. The switch between transcription and replication is poorly understood but is likely regulated to ensure the production of viral proteins and vRNPs in the correct stoichiometric proportions.

Despite the importance of understanding IAV transcription and replication dynamics, progress has been impeded by the lack of available tools to study infection processes at single -cell resolution. Traditional bulk analysis methods, such as RT-qPCR and RNA-seq, provide averaged measurements across populations of cells, masking the inherent heterogeneity of viral infection. Methods capable of visualizing individual RNA molecules within cells allow the exploration of cellular heterogeneity, amongst which padlock probe (PLP) based technologies show particular promise for following viral infections. PLP methodologies employ DNA oligonucleotide probes containing 5’ and 3’ arms designed for specific hybridization to their target and a non-hybridized programmable loop region (**Fig. 1a**) (3–6). The loop usually contains specific binding sites for different fluorophores (7–11), but can also contain molecular ‘barcodes’ that can be read out by *in-situ* (6,12–14) or *ex-situ* sequencing (14). Following hybridization to their target, PLPs are ligated into circular molecules for subsequent rolling circle amplification (RCA). RCA produces a nanoball rolling circle product (RCP) containing thousands of concatemeric antisense copies of the circularized PLP, enabling the detection of single RNA molecules (13). Due to the stringency afforded by the ligation step (15), PLPs exhibit great specificity, meaning that highly similar RNA molecules can be readily distinguished. However, this specificity comes at the expense of sensitivity. This is generally attributed to both the inefficient binding of the PLP to its target and the inefficient conversion of RNA molecules into cDNA during *in situ* reverse transcription, which is the classical target of PLPs (6). Recently, direct ligation of the probes on the RNA target has been reported to enhance the overall sensitivity of PLPs (6). This utilizes PBCV-1 DNA ligase, which can perform ligation of single-stranded DNA when splinted by a complementary RNA strand (16). As PLPs targeting distinct regions can contain the same loop region, distinct probes can be programmed with the same fluorophore binding site or barcode. This offers a potential way to further increase sensitivity by increasing the chance that a target is bound by at least one PLP.

**Figure 1.**
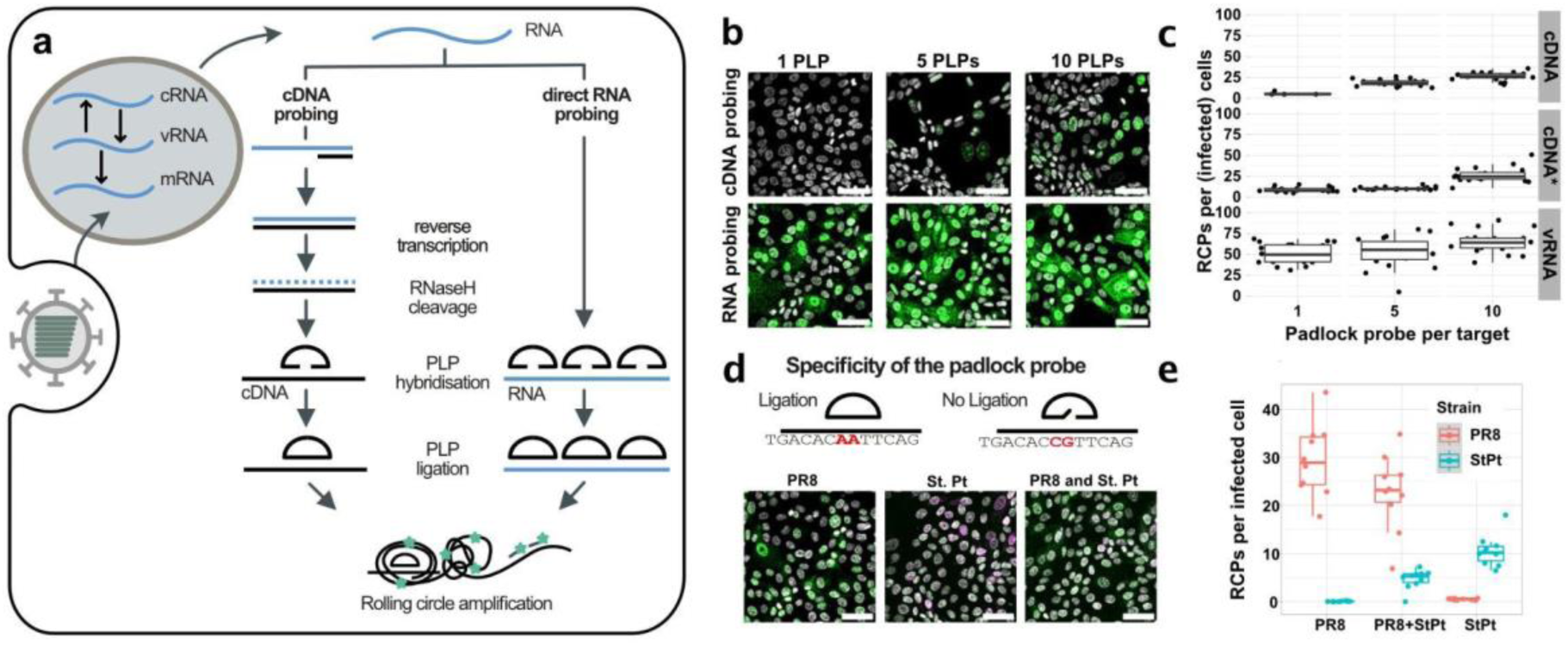
Direct RNA padlock probing enables highly sensitive and strain-specific detection of Influenza A virus genomes. (a) Schematic comparison of classical cDNA padlock probing and direct-RNA padlock probing. After hybridization to the targets (cDNA or RNA), the padlock probes are ligated, followed by rolling circle amplification (RCA) to generate concatemers. The loop region of the padlock probe contains the detection probe sequence and after RCA, the rolling circle products (RCPs) are hybridized with a detection probe and detected as spots. (b) Sensitivity comparison between classical padlock probing and direct RNA padlock probing. MDCK cells were infected with 1 MOI and samples were probed for the PB1 segment (either direct RNA or cDNA) after 6 hpi (scale bar = 50 µm). (c) Graph showing the average RCPs detected per infected cells in case of direct RNA probing and cDNA probing. Y-axis is limited to the range 0-100, one FOV with 196 RCPs per infected cell in 5PLP vRNA is not shown. cDNA* is sample padlock probed with additional primer to prime the RCA reaction (d) Specificity assessment of the direct RNA padlock probing strategy using two closely related H1N1IAV strains i.e. PR8 (in green) and St.Petersburg (StPt) (in magenta). Samples were infected with PR8 only, PR8 and StPt (co-infection), or StPt only and all the samples were padlock probed for both strains probe set. (scale bar = 50 µm). (e) Quantification demonstrating the specificity of the direct RNA padlock probing strategy. RCP spots were detected for both padlock probes (PLP) sets only in the case of co-infection, confirming strain-specific detection.

In this study, we introduce multiple direct-RNA assisted padlock probes in combination with *in-situ* sequencing (mudRapp-seq) to enable the sensitive and specific localization of IAV RNAs in infected cells. We show that direct RNA targeting PLPs offers a three-fold higher detection efficiency compared to PLPs targeting cDNA molecules. We also demonstrate that employment of five to six PLPs per target improves detection efficiency without compromising specificity. The inclusion of barcodes within the programmable loop region of the PLP coupled with *in-situ* sequencing allows for highly multiplexed detection of target RNAs. We apply mudRapp-seq to simultaneously visualizing 16 IAV mRNAs and vRNAs in infected cells, and to distinguish between IAV vRNAs of highly similar strains during co-infection. With the implementation of cellular segmentation, mudRapp-seq identifies the expression of the M segment mRNA as a potential factor in regulating the switch between transcription and replication. Thus, mudRapp-seq enables the high fidelity analysis of multiple RNA species within individual cells, enhancing our understanding of transcriptional heterogeneity and the spatial localization of gene expression changes in situ.

## Material and Methods

### Cell culture and virus propagation

Regular maintenance and passage of cells were performed in standard cell culture dishes. Madin-Darby canine kidney cells (MDCK) and Human Embryonic Kidney 293 T (HEK 293T) cells were cultured in Dulbecco’s Modified Eagle Medium (DMEM) (supplemented with 10% heat-inactivated fetal bovine serum and 100 U/mL penicillin-streptomycin, Gibco, Life Technologies) in a 5% CO2 humidified atmosphere at 37 °C. Laboratory-adapted A/Puerto Rico/8/1934 (PR8) strain was rescued using a reverse genetic approach. 2 mL of trypsinized HEK cells were seeded in 6 well plates at a density of 1×10^6^ cells/mL in a supplemented DMEM medium. A total of 600 ng of each plasmid segment were added in PEI (1 mg/mL) with the ratio 1:12 and mixed by vortexing, followed by the addition of 500 µL of opti-MEM (Gibco, Life Technologies). The DMEM medium was removed from the well, and fresh opti-MEM was added followed by the addition of the transfection mix dropwise to the cell. The cells were incubated at 37 °C for 6 h. Afterwards the medium was changed, and fresh opti-MEM was added to the cells which were then incubated for a total of 48 h. Following this incubation, the supernatant was collected and added to MDCK cells for viral propagation. After one hour of incubation, the supernatant was again removed and replaced with fresh opti-MEM medium with Tosyl phenylalanyl chloromethyl ketone (TPCK). The cells were then incubated for an additional 32-48 h. The supernatant was collected and centrifuged at 15000 rpm for 15 min, aliquoted and stored at –80 °C.

### Viral infection

MDCK cells were grown in an 18-well chamber slide (µ-Slide 18 well glass bottom, Ibidi) for 6-10 h in DMEM medium at 37 °C and then washed with 1x PBS with Mg^2+^ and Ca^2+^ supplemented with 0.2% BSA. Viral stocks were diluted to the appropriate multiplicity of infection (MOI) and cells were infected for 1 h at 37 °C. Following infection, the cells were washed with 1x PBS and incubated in opti-MEM media containing TPCK to further promote viral replication.

### Fixation

After the desired incubation period, the cells were washed with 1x PBS and fixed with 4% formaldehyde (Methanol free formaldehyde, Pierce, Life Technologies) for 10-15 minutes. Next, samples were permeabilized with 70% ethanol for 30 min at room temperature or overnight at 4°C. Before further processing, the samples were washed twice with 1x PBS containing tween-20 (1xPBS-T).

### Padlock probe phosphorylation

All Padlock probes were phosphorylated before usage. A mixture containing 1 mM ATP, 0.2 U/µL T4 polynucleotide kinase (PNK) (NEB) and 1x PNK buffer was added to each padlock probe (2 µM) and incubated at 37°C for 30 min and 65°C for 10 min. The phosphorylated probes were stored at -20°C until further usage.

### Padlock probing

Fixed and permeabilized samples were first washed with 1x-PBS. Padlock probing was carried out either on cDNA or directly on RNA.

For cDNA padlock probing, cells were washed with 1x-PBS containing 0.05% Tween 20 (PBST), for 10 min, and washed twice with 1x PBS. cDNA synthesis was then performed overnight at 37 °C using 5 U/µL RevertAid H minus reverse transcriptase (EP0752, Thermo Fisher Scientific) in 1x RevertAid RT buffer with 0.25 mM dNTPs, 0.2 mg/mL BSA, 5 µM mixture of randoms 15 mer primers and target-specific primers for PB1 segment (TTACAGGTACACGTACCGATGCC, TCTAAATAGAAACCAACCTGCTGCAAC, AAAGCAGGCAAACCATTTGAATGGATG, TCACATATATGACCAGAAATCAGCCCG) (Integrated DNA Technology, IDT) and 0.8 U/µL Rnasin. After overnight reverse transcription, the sample was washed three times with 1x PBS-T. cDNAs were fixed with 4% (w/v) formaldehyde for 15 min followed by three times washing with 1x PBS-T and one time with 1x PBS. PLPs with a concentration of 0.1 µM of each were annealed and ligated simultaneously using 0.5 U/mL Ampligase (Lucigen, Biozym) in 1x Ampligase reaction buffer containing 20% formamide, 0.2 mg/mL BSA, 50 mM KCl, and 0.4 U/mL RNase H for 30 min at 37 °C followed by 45 min at 45 °C.

For RNA padlock probing, the padlock probe was diluted in hybridization buffer (6x SSC and 10 % formamide) to a final concentration of 0.1 µM along with 0.2 µM of RCA primer (For mRNA and vRNA dynamic experiment; vRNA segments: GTGCGAGAGCGTCGGTATTAAGC, mRNA GCGGCTCCACTAAATAGACGG, cDNA and direct RNA as well as for specificity and sensitivity experiments: GCGGCTCCACTAAATAGACGG. The mixture was then added to the samples and hybridized for 1 h at 37 °C. The sample was washed with the hybridization buffer for 5-10 min, followed by washing with 1x PBS-T and 1x splint-R ligase buffer for 5 min. The probes were ligated with splint-R ligase with a ligation mix containing 1x splint-R ligase buffer and 210 nM splint-R ligase (M0375S, New England Biolabs, (NEB)) for 1 h at room temperature.

After ligation, samples were washed three times with 1xPBST for 10 min followed by a wash with 1x PBS. The samples were washed with 1x Phi-29 DNA polymerase buffer for 5 min and then incubated with RCA mix (1 U/µL Phi-29 DNA polymerase (NEB), 250 µM dNTPs 200 µg/ml BSA, 40 µM amino-ally dUTP (NU-803S, Jena Bioscience), 5 % glycerol) in 1x Phi-29 DNA polymerase buffer for 12-16 h at room temperature. After RCA, the rolling circle products (RCPs) were cross-linked 50 mM BS(PEG)9 (PEGylated bis(sulfosuccinimidyl)suberate) in 1x PBS (Thermo scientific) for 1 h, followed by neutralization with 1 M Tris-HCl at pH 8.0 for 30 min at room temperature, and washed three times with 1x PBS-T for 5 min.

### RCP detection

RCPs were detected either directly with a fluorescently labeled primer (IDT) or using sequencing by synthesis. In both cases, samples were first washed three times with 1x PBS-T, one time with 1x PBS, and finally rinsed with a hybridization buffer.

For direct detection of RCPs, samples were hybridized with 500 nM of fluorescently labeled primer (/FluorT/GCGTCTATTTAGTGGAGCCGC) in hybridization buffer (6x SSC buffer and 10 % Formamide).

For sequencing by synthesis, samples were then hybridized with 500 nM of sequencing primer in hybridization buffer for 30 min at room temperature, followed by a wash with 100 µL hybridization buffer and three washes with 1x PBS-T (for the mRNA and vRNA dynamic experiment; vRNA segments: GTGCGAGAGCGTCGGTATTAAGC, mRNA GCGGCTCCACTAAATAGACGG, for specificity experiment: GCGGCTCCACTAAATAGACGG). Sequencing-by-synthesis was performed using the reagents from an Illumina MiSeq 500 cycle Nano kit (MS-103-1003 Illumina). In the first step, the samples were washed with incorporation buffer (Nano kit PR2) and then incubated for 13 minutes at 50 °C in incorporation mix (Nano kit reagent 1). Subsequently, the samples were thoroughly washed three times with PR2 buffer at room temperature followed by three washes at 50 °C. Next, the samples were placed in a solution of 200 ng/mL DAPI in 2x SSC buffer to counterstain the nucleus, followed by a further two washes with incorporation buffer. After that imaging mix (Nano kit reagent 2) was added and the samples were imaged. Following imaging, dye terminators were removed by incubating the samples with Illumina cleavage mix (Nano kit reagent 4) at 50 °C for 15 minutes, followed by thorough washing with PR2 to remove any residual cleavage mix, allowing for the successful initiation of the next round of incorporation.

### Image acquisition

All images were acquired using an epifluorescence microscope (Leica) with automated XYZ stage control and hardware autofocus. An LED light engine was used for fluorescence illumination and Leica-DFC9000GT-VSC11903 camera. Fluorescently labeled primers were imaged using FluorT labeled primers obtained from integrated DNA technology (IDT). In-situ sequencing cycles were imaged using a HC PL APO 63x/1.40 oil objective with the following filters and exposure times for each base: G (excitation 506/21 nm, emission 535/70 nm, dichroic 523 nm, 100 ms); T (excitation 578/24 nm, emission 642/80 nm, dichroic 598 nm, 50 ms); A (excitation 638/31 nm, emission 642/80 nm, dichroic 660 nm, 100 ms); C (excitation 730/40 nm, emission 810/80 nm, dichroic 740 nm, 200 ms). The nuclear DNA was stained with DAPI and imaged at excitation 391/31 nm, emission 460/80 nm, dichroic 415 nm, 50 ms.

### Image analysis

Images were computationally cleared (ICC) and z-stacks maximum intensity projected using the Leica Application Suite X version 3.8.1 (LAS X 3.8.1). The images of the different rounds of incorporation were registered and cropped to the common region by a translation learned from the maximum intensity projection of the fluorescent images across channels (A,C,G,T). Background removal with a White TopHat filter and bleed-through correction with LinearUnmixing on the sequencing channels were used, followed by histogram matching, clipping with re-scaling, bandpass and median filtering (using a modified version of starfish 0.2.2, https://github.com/BioMeDS/starfish/tree/947472f79ca5206bc9075c7bb45bb28425b3f105). After RCP spot detection, barcodes were decoded according to the mean intensity of each sequencing channel within the spot region. Nuclei and cells were segmented using separate cellpose 2.2.3 models (17). Cell segmentation masks were manually curated in napari 0.4.17(https://github.com/napari/napari). All analysis code is openly available at https://github.com/BioMeDS/mudRapp-seq.

### Statistical analysis

A linear model for the relationship between segment wise mRNA abundance and total vRNA abundance was fitted to the counts of all cells with at least 40 mRNA RCP spots from at least 6 distinct mRNA segments. The mRNA abundances were added as independent predictors (no interaction terms) using the lm function (18) of R version 4.4.1 (https://www.R-project.org).

### Nano-DMS-MaP probing

#### Virus infection, DMS probing and RNA extraction

MDCK cells were cultured in 6 well plates and infected with 1 MOI of PR8. The infected cells were fixed and permeabilized at 6 hpi. The padlock probes were then hybridized in a hybridization buffer for 1 h at 37 °C and washed three times, followed by ligation of the PLPs with splint R ligase (as described above). For DMS-probing, 1 mL of a 32 mM DMS solution diluted in DPBS 1X (Gibco) was added to the cells and incubated for 6 min at 37 °C. The reaction was immediately quenched by transferring cells on ice and adding 10 µL of β-mercaptoethanol 14 M (Sigma-Aldrich). Cells were washed three times with DPBS 1X, scrapped and proteinase K treated in lysis buffer (20 mM Tris-HCl pH 7.5, 150 mM KCl, 1 mM MgCl2, 1 mM DTT, 1 mM PMSF, 0.2 % Triton x100, 200 U/mL RnaseIn and Proteinase K 80 U/mL) at 37 °C for 1 h. 1 mL of TRI reagent (Sigma Aldrich) was added and incubated for 5 min at room temperature and the solution was transferred to 2 mL tubes. RNA was extracted according to the manufacturer’s instructions.

#### Reverse transcription and PCR amplification of HA and NA segments

HA and NA vRNA segments were reverse transcribed using the primer HA_vRNA_RT (AGCAAAAGCAGGGGAAAATAAAAACAACCAAAATG) for HA, and NA_vRNA_RT (AGCAAAAGCAGGAGTTTAAAATGAATCCAAATCAG) for NA. 1 µg of extracted RNAs were mixed with 12.5 nmol dNTPs and 5 pmol of RT primer in 9 µL of RNase-free H_2_O and denatured for 5 min at 65 °C. Samples were placed on ice for 2 min and reverse transcription was performed in a final volume of 25 µL containing 50 mM Tris–HCl pH 8.3, 200 mM KCl, 5 mM DTT, 20% glycerol, 1 mM MnCl2, 0.32 U/µL RNaseIn and 1.6 U/µL Marathon-RT (19) for 8 h at 42 °C.

Viral RNA species were differentially amplified using the primers PCR_HA_Fw (AGCAAAAGCAGGGGAAAATAAAAACAACCAAAATG) and PCR_HA_Rv (AGTAGAAACAAGGGTGTTTTTCCTCATATTTCTG) for the full-length HA vRNA segment, and PCR_NA_Fw (AGCAAAAGCAGGAGTTTAAAATGAATCCAAATCAG) and PCR_NA_Rv (AGTAGAAACAAGGAGTTTTTTGAACAGACTACTTG) for NA vRNA. 5 µL of 1/5 diluted cDNA was used as template in 50 µL PCR reaction containing 0.05 U/µL PrimeSTAR GXL polymerase (Takara Bioscience), 250 nM of each Fw and Rv primers, 200 µM of each dNTP and 1× PrimerSTAR GXL buffer in a total volume of 50 µL. Cycling conditions were initial denaturation for 2 min at 98 °C, followed by 25 cycles for 15 s at 98 °C, 20 s at 55 °C, 2 min at 68 °C, followed by a final extension for 7 min at 68 °C. Amplicon quality was checked on a 1 % agarose gel post stained in ethidium bromide solution.

#### Nanopore sequencing and analysis

DNA amplicons were purified with Mag-Bind® TotalPure NGS (Omega) using a volume of beads of 0.7× the sample volume and following the manufacturer’s instructions. Next, 80 ng of purified amplicon DNA was diluted with nuclease-free water in 5 µL. Simultaneous dA-tailing and 5′-phosphorylation was performed by addition of 0.7 µL NEBNext End-Repair Buffer and 0.3 µL NEBNext End-Repair enzymes (NEB), followed by an incubation at room temperature for 5 min and an inactivation step at 65 °C for 5 min. Barcodes of the kit SQK-NBD114.96 (Oxford Nanopore Technologies) were then ligated in a 10 µL reaction containing 3.5 µL end-repaired DNA, 1.5 µL barcode (SQK-NBD114.96) and 5 µL NEB Blunt/TA Ligase Master Mix for 20 min at room temperature. The ligation was terminated by addition of 1 µL EDTA (SQK-NBD114.96). Samples were pooled and purified with 0.4× Ampure XP beads (SQK-NBD114.96), and washed twice with short fragment buffer (SFB, SQK-NBD114.96). After elution of the pooled barcoded DNA in 25 µL H_2_O, dsDNA was quantified using AccuClear® Ultra High Sensitivity dsDNA Quantitation Solution (Biotium). Native adapter (NA) was then ligated in a 50 µL reaction containing 200 fmol of barcoded DNA, 10 µL NEBNext Quick Ligation Reaction Buffer (NEB B6058S), 5 µL NA adapter (SQK-NBD114.96) and 5 µL NEB T4 DNA ligase high concentration (NEB T2020M), incubated for 20 min at room temperature. A final purification was performed with 0.4× Ampure XP beads and SFB buffer (SQK-NBD114.96), and the library was eluted in 15 µL of Elution buffer. After quantification with AccuClear, 75 fmol of the final library was sequenced on a PromethION R10.4.1 flow cell (FLO-PRO114M, Oxford Nanopore Technologies) on a P2 Solo sequencer (Oxford Nanopore Technologies) using MiniKnow acquisition software (Oxford Nanopore Technologies) v.21.11.8. The resulting data was basecalled and demultiplexed using dorado (Oxford Nanopore Technologies) v0.5.3 with the dna_r10.4.1_e8.2_400bps_sup@v4.2.0 model.

DMS-MaP sequencing data was aligned to the reference sequence using LAST v1548 (20), with the parameters ‘-T1 -Qkeep -m20 -p. The mismatch matrix was generated using the last-train tool for each package. Further analysis was carried out using the rf-count and rf-norm modules of RNA Framework package v2.7.2 (21). rfcount parameters were only-mut ‘G>Y;A>B;C>D;T>V’ -m -nd -ni -q 16 -eq 10’. rf-norm parameters were ‘-rb AC -sm 3 -nm 2 -- max-untreated-mut 0.05 --max-mutation-rate 0.5 --norm-independent’, meaning that DMS reactivities were calculated by subtracting background mutations in the untreated sample and normalized using 90% Winsorizing (22).

#### Western blot

MDCK cells were infected with PR8 at an MOI of 5. At the indicated timepoints, whole cell extracts were harvested using Nonidet P-40 lysis buffer (20 mM Tris HCl, 137 mM NaCl, 10% Glycerol, 1% NP-40 and 2 mM EDTA). Protein samples were separated on 15% polyacrylamide gels (Bio-Rad) and transferred onto nitrocellulose membranes (Bio-Rad). Membranes were blocked with 5% (wt/vol) skim milk in DPBS x1 for 1 h at room temperature (RT) and incubated overnight at 4°C with primary antibodies: anti-M1 (1 μg/mL; GeneTex GTX125928), anti-M2 (1 μg/mL; Santa Cruz sc-32238), anti-NP (1 μg/mL; BEI Resources NR-43899), or anti-actin (1 μg/mL; Invitrogen MA611869) in 5% (wt/vol) skim milk. After washing, membranes were incubated for 1 h at 25 °C with secondary antibodies: ECL-linked anti-mouse (1:1000; Cytiva NA931V) or ECL-linked anti-rabbit (1:1000; Cytiva NA974V). Protein detection was performed using Immobilon Western Chemiluminescent HRP Substrate (Millipore) according to the manufacturer’s instructions.

#### Immunofluorescence staining

MDCK cells were grown on coverslips and infected with PR8 at an MOI of 0.3 or 1. At the indicated time points, cells were rinsed twice with PBS and fixed for 10 min in Histofix (4% formaldehyde, Carl Roth) at 4°C. Following fixation, cells were rinsed twice with PBS, permeabilized with 0.5% NP-40 in PBS for 10 min at room temperature (RT), and subsequently blocked in 5% bovine serum albumin (BSA) in PBS for 30 min. Cells were then incubated for 2 h at RT with primary antibodies: anti-M1 (1:500; GeneTex GTX125928), anti-M2 (1:100; Santa Cruz sc-32238), or anti-NP (1:1000; BEI Resources NR-43899). After primary antibody incubation, cells were washed and incubated for 1 h at RT with species-appropriate secondary antibodies: CF568-conjugated anti-rabbit (1:1000; Sigma SAB4600076), CF488-conjugated anti-mouse (1:1000; Sigma SAB4600035), or Alexa Fluor 555 (AF555)-conjugated anti-mouse (1:1000; Invitrogen A21425). Nuclei were counterstained with Hoechst dye at a final dilution of 1:10,000.

#### Effect of RNP and M pre-expression by RT-qPCR

For testing the effect of pre-expression of RNP (NP, PA, PB1, PB2) and M, reverse transfection of 600 ng total DNA in HEK293T cells using PEI, as described above. 24–48 h post-transfection, the culture medium was removed, and the cells were washed once with DPBS x1. Subsequently, HEK293T cells were infected with PR8 virus at MOI 5 without the addition of TPCK-trypsin. Viral adsorption was carried out at 37°C for 1 h, with gentle shaking every 15 min to enhance infection efficiency. Following adsorption, the viral inoculum was removed, and the medium was replaced with opti-MEM containing CHX at 0, 50, or 100 µg/mL. Samples were collected at 4 and 6 hpi.

RNA was extracted using the Trizol method, followed by a 270-fold dilution of the RNA and an additional 32-fold dilution for 18S rRNA. Reverse transcription was performed using specific primers (HA vRNA: agcaaaagcaggggaaaataaaaacaaccaaaatg, HA mRNA: TTTTTTTTT TTTTTTTTTTTV N, 18S rRNA: TAATGATCCTTCCGCAGGTTCACCTAC), and the resulting cDNA was diluted 10-fold before quantitative PCR (qPCR). qPCR was conducted using the PowerUp SYBR Green system, targeting vRNA (HA), vmRNA, and 18S rRNA using following primers (HA vRNA and mRNA are the same: GCTCATGGCCCAACCACAACACAAACG, GGACTTCTTTCCCTTTTTTGTTCACATAAG, and for 18S rRNA: CGAGGATCCATTGGAGGGC, CCGCTCCCAAGATCCAACT, with data analyzed using the ΔΔCt method.

## Results

### Multiple direct RNA-assisted padlock probing allows the sensitive and specific detection of RNA *in situ*

Padlock probes (PLPs) are widely used for the detection of RNA molecules in cells (23). In conventional PLP methodologies, the RNA target is first reverse transcribed into cDNA and then treated with RNase H to free the cDNA. PLPs are then hybridized to the cDNA target and ligated with a DNA ligase to generate a circular template for RCA (4). Direct-RNA padlock probing represents an alternative approach wherein the PLP is directly hybridized to the target RNA itself and then circularized into a template for RCA using PBCV-1 ligase (Paramecium bursaria Chlorella virus 1 ligase), an enzyme capable of ligating single-stranded DNA molecules when splinted by a complementary RNA strand (**Fig. 1a**). Whilst direct RNA padlock probing is reported to improve RNA detection efficiency, viral RNAs are heavily complexed with viral proteins potentially preventing their accessibility to direct RNA padlock probes. We therefore set out to assess whether viral RNAs can be detected using this new approach. For this, we infected MDCK cells at 1 multiplicity of infection (MOI) with the laboratory-adapted strain H1N1, A/Puerto Rico/8/34 (PR8). Infected cells were fixed and permeabilized 6-h post-infection (hpi), and vRNA for the PB1 segment was visualized using both cDNA and direct RNA methodologies in parallel (**Fig. 1b**). With a single PLP targeting PB1 vRNA, we found a mean of 6.0 RCPs per infected cell when RNAs were detected through a cDNA intermediate (**Table 1**). We detected these RCP spots in only four cells, and they were predominantly found in the nucleus. In contrast, direct RNA probing led to the mean detection of 50.2 RCPs per infected cell with 98% of cells having at least 5 RCPs, as expected when MDCK cells are infected with an MOI of 1 (**Table 1**). These results demonstrate that direct RNA probing enhances sensitivity, even on highly proteinated viral RNAs.

**Table 1:** Comparison of cDNA probing and direct RNA probing efficiencies. Direct RNA probing is 2-3 times more sensitive than cDNA probing. Detection efficiency increases with an increasing number of padlock probes per target.

|  | cDNA |  |  | cDNA with RCA primer |  |  | direct RNA |  |  |
| --- | --- | --- | --- | --- | --- | --- | --- | --- | --- |
|  | Percentage of RCP spots in the nucleus | Percentage of infected cells | Mean RCP spots per infected cells | Percentage of RCP spots in the nucleus | Percentage of infected cells | Mean RCP spots per infected cells | Percentage of RCP spots in the nucleus | Percentage of infected cells | Mean RCP spots per infected cells |
| **1 PLP** | 0.827 | 0.020 | 6.0 | 0.851 | 0.120 | 9.5 | 0.676 | 0.980 | 50.2 |
| **5 PLPs** | 0.835 | 0.457 | 18.5 | 0.858 | 0.368 | 10.7 | 0.585 | 0.953 | 57.3 |
| **10 PLPs** | 0.661 | 0.529 | 27.0 | 0.777 | 0.656 | 26.1 | 0.575 | 0.953 | 64.4 |

To further optimize detection efficiency, we performed the same experiments with 5 or 10 PLPs distributed evenly across the entire length of segments. These PLPs target the same vRNA without spatial overlap (**Fig. 1b**, **1c**, and **Table 1**). For the cDNA experiment, increasing the number of PLPs boosted detection efficiency with 18.5 and 27.0 RCPs per infected cell seen with 5 and 10 PLPs, respectively (**Table 1**). Now, 46% and 53% of cells contained at least 5 RCPs for cDNA PLP at 5 and 10 PLP, respectively (**Table 1**). As before, the RCPs were mainly localized to the nucleus where viral RNAs and their replication intermediates are concentrated. We also tested a cDNA PLP with an additional primer (cDNA*) to prime the RCA reaction, but this did not lead to an improvement in RNA detection (**Fig. 1b**, **1c and Supplementary Fig. 1**). For direct RNA padlock probes, increasing the number of PLPs led to the detection of marginally more RCPs (64.4 mean per infected cell with 10 PLPs), although the signal crowding in the nucleus may have led to an underestimation of the true signal (**Table 1**). Under all direct RNA PLP conditions, extensive localization of the PB1 vRNA was also seen in the cytoplasm where assembly intermediates localize to Rab11-dependent vesicles (24,25) (**Fig. 1b**). This indicates that direct RNA PLPs can be more effective at imaging low abundance RNAs than cDNA PLPs due to their improved detection efficiencies.

To assess the specificity of our approach, we performed the same experiments on uninfected cells. We observed almost no signal for both cDNA and direct RNA PLPs, demonstrating low background and high specificity (**Supplementary Fig. 1**). For a more rigorous evaluation, we designed additional PLPs against another H1N1 strain, A/St.Petersburg/8/2006 (StPt), which is highly similar to the H1N1 vaccine strain PR8 used previously. These two strains share 80-90% sequence identity, which makes it challenging to design single-molecule fluorescence in situ hybridization (smFISH) probes that specifically hybridize to one strain but not the other. However, PLP-based methodologies can faithfully discriminate even single nucleotide polymorphisms due to the ability of DNA ligases to discriminate mismatches (26). Exploiting this property, we designed 10 PLPs against HA vRNA segment containing at least one nucleotide mismatch at the ligation junction (**Fig. 1d**). We included both probe sets in mock, single- and co-infected cells. As before, mock-infected cells showed a very low background with both PLP sets (**Supplementary Fig. 1**). Importantly, we only observed signals when the PLP set matched the strain used for the infection, or when cells were co-infected (**Fig. 1e**). These results collectively demonstrate that mudRapp provides excellent sensitivity without compromising specificity.

### PLP efficiency is driven by target site availability, binding, and ligation efficiency

To elucidate factors influencing improved PLP efficiency with multiple PLPs, we probed for HA and NA vRNAs using individual PLPs (**Fig. 2a-b** and **Supplementary Fig. 2a-b**). Each PLP was designed to hybridize to the target with approximately the same binding energy using binding arms of approximately the same length (35-40 nt) (**Supplementary Data 1**). Imaging revealed substantial differences in detection efficiency, with certain PLPs showing almost no RCPs, whereas others detected 40-50 RCPs per cell (**Fig. 2c, Supplementary Fig. 2c,** and **Supplementary Data 1**). As before, we consistently observed very low background in mock-infected cells confirming that the individual PLPs are highly specific (**Fig. 2d** and **Supplementary Fig. 2d**). To determine the optimal number of PLPs, we titrated increasing numbers of individual probes per target. We found that more PLPs generally led to more RCPs, with detection efficiency reaching saturation when sample were probed with 5-6 PLPs (**Fig. 2e-g** and **Supplementary Fig. 2e-g**).

**Figure 2.**
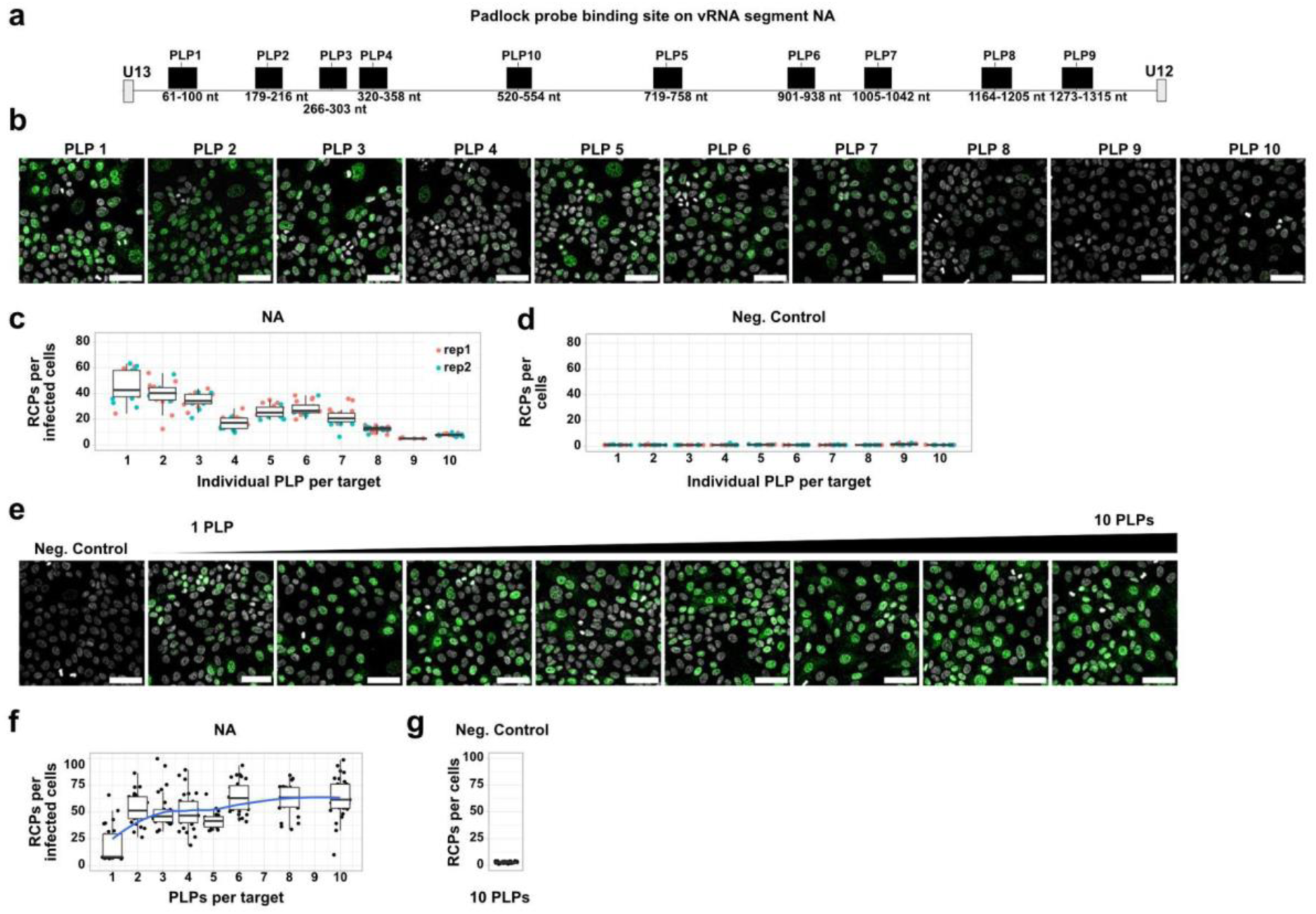
Multiple-direct RNA padlock probing improves the detection efficiency of the RNA targets. (a) Schematic representation of padlock probe (PLP) binding sites on the NA segment of IAV strain PR8 vRNA. (b) Images demonstrating heterogeneity in PLP binding across different regions of the NA segment (scale bar = 50 µm). (c) Binding efficiency of individual PLPs for the NA segment. (d) Quantification of unspecific binding of the individual padlock probe. Negative control was performed on non-infected MDCK cells. (e) Probing PR8 infected MDCK cells (1 MOI) after 6 hpi with increasing number of padlock probes. (f) Graph showing the increase in target detection efficiency as a function of the number of padlock probes per target. The blue line represents smoothed data obtained through local polynomial regression fitting. (g) Negative control with all 10 NA PLPs on uninfected cells.

We hypothesized that the variable performance of individual PLPs results from inefficient probe binding, likely due to RNA secondary structures limiting accessibility to certain target regions. To test this, we employed long read RNA structural probing by nanopore dimethyl sulfate mutational profiling (Nano-DMS-MaP) (27,28). DMS is a cell-permeable chemical that methylates adenines (A) and cytosines (C) in single-stranded RNA regions but does not react with bases engaged in RNA secondary structure or base-paired to PLPs (**Fig. 3a**). These DMS modifications cause nucleotide misincorporations during reverse transcription, which can be detected as mutations upon nanopore sequencing. Using this approach, we obtained comprehensive DMS reactivity profiles across the full length of the NA and HA vRNAs (**Supplementary Data 2** and **3**). We included control samples without PLPs to inform on accessible and inaccessible region within the native vRNAs. Samples with PLPs were used to identify PLP hybridisation sites by comparison with the native RNA structure. For NA, the DMS reactivity profiles from two biological replicates were highly reproducible (Pearson’s r > 0.97). When comparing windowed profiles (window size=20) in the absence and presence of PLPs we observed clear protections from DMS at the predicted target sites, and comparatively few changes outside of the target site. Together with the lack of detectable RCPs in mock infected cells (**Fig. 2d** and **Fig. 2g**), these data indicate highly specific binding with little evidence of off-target binding (**Fig. 3b**).

**Figure 3.**
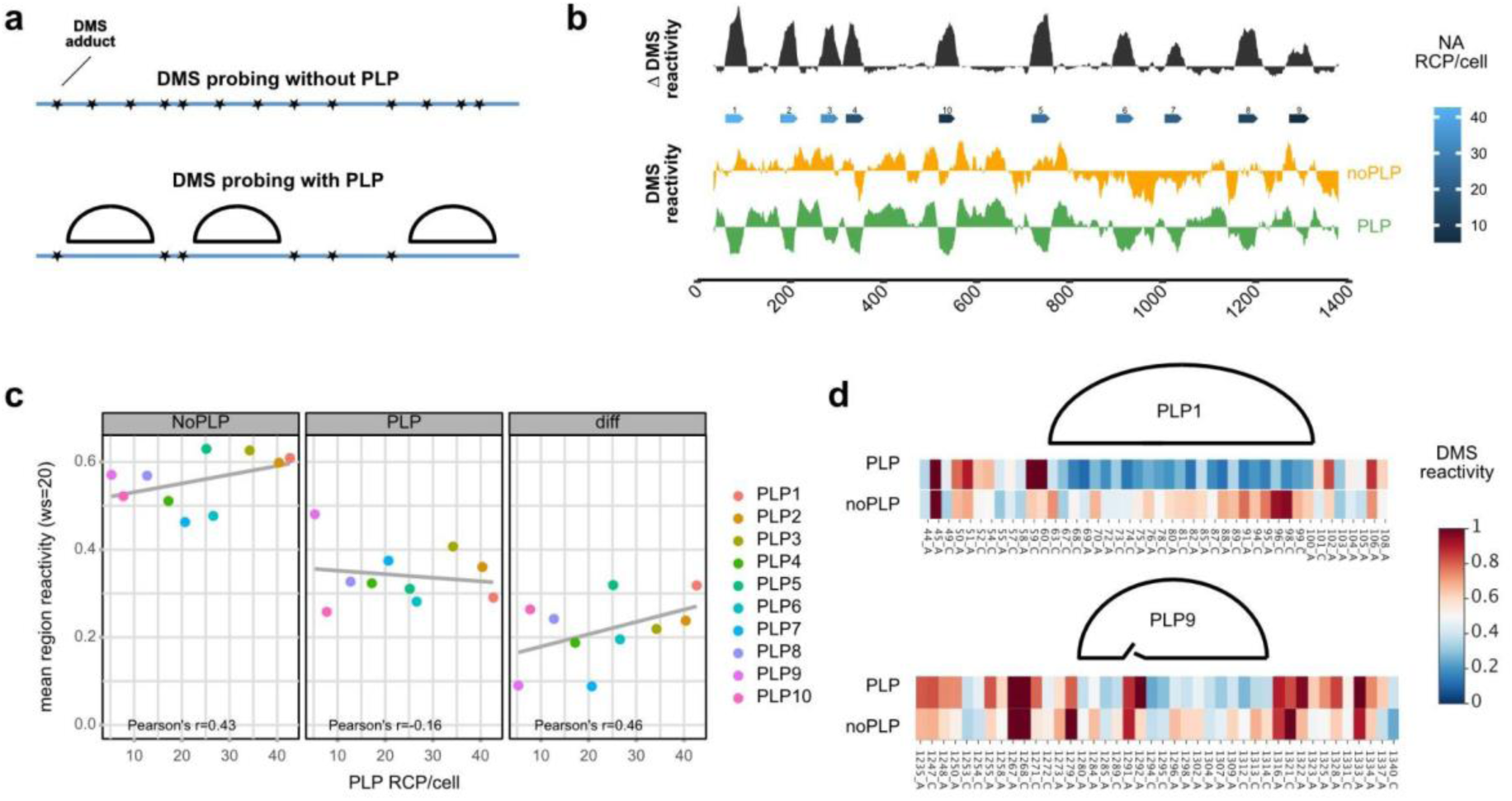
DMS probing reveals the impact of RNA structure on padlock probe binding efficiency. (a) Schematic illustration of DMS (dimethyl sulfate) probing principle: single stranded RNA regions are reactive to DMS, base-paired RNA and the region hybridized with padlock probe will be totally protected from DMS modification. (b) DMS reactivity profile with and without padlock probe binding along the NA viral RNA segment. Individual PLPs are displayed as blue arrows, coloured by binding efficiency as measured by RCP/cell. Illustration generated with genomes. (c) Correlation analysis of DMS reactivity at target sites with and without padlock probe binding. The scatter plot shows the relationship between DMS reactivity in both conditions, with the difference highlighting structural changes upon probe binding. (d) Heat map plot of the high binding region (PLP1) and low binding region (PLP9) on the NA segment. PLP9 binding site has high DMS reactivity at the ligation junction.

Notably, NA detection efficiency was positively correlated with DMS reactivity at the target site in the absence of PLPs (Pearson’s r=0.43), suggesting that PLP efficiency is enhanced when the target site is unstructured (**Fig. 3c**). Additionally, we observed a negative correlation between detection efficiency and DMS reactivity in the presence of PLPs (Pearson’s r=-0.16), consistent with the expectation that PLP binding shields the target site from DMS modification (**Fig. 3c**). Correspondingly, PLP efficiency was best correlated with the mean difference in DMS reactivity in the presence and absence of PLPs (Pearson’s r=0.46), indicating that both initial accessibility and subsequent binding strength contribute to effective detection (**Fig. 3c**).

Interestingly, for NA more pronounced correlations were found when DMS reactivity windows were centered at the ligation junction indicating that certain PLPs may bind correctly to their target without being efficiently ligated, likely due to sequence-dependent ligase preferences (16) (**Supplementary Fig. 3**). For instance, the worst performing PLP, NA_PLP9, exhibited DMS protection across the target binding site but showed very high reactivity to DMS at nucleotides A1291-A1292, precisely at the ligation junction (**Fig. 3d**). Similar DMS reactivity profiles and correlations to number of RCPs were obtained for the HA segment, except for HA_PLP6 and HA_PLP8, which showed very high PLP efficiencies despite highly structured RNA at the target site (**Supplementary Fig. 4a-b**). These two PLPs presented the lowest DMS reactivities at the target site in the presence of the PLPs, indicating that they were able to bind with very high efficiencies despite a structured target site. For HA, however, we did not observe a better correlation between DMS reactivities and the number of RCPs per cell when the ligation junction was considered (**Supplementary Fig. 4c**).

Altogether, these data demonstrate that multiple direct RNA PLP can increase detection efficiency by overcoming inefficiencies in PLP binding and ligation, without compromising its exquisite target specificity. Moreover, open RNA structure at the target site is a predictor, but not a prerequisite, of PLP efficiency.

### Multiplexed RNA detection using barcodes read out by *in-situ* sequencing

Padlock probing is ideally suited for multiplexing because the loop region of the PLP can contain arbitrary sequences that can bind distinct fluorescently labeled probe (**Fig. 4a**). However, the spectral overlap of common dyes limits the number of simultaneously detectable transcripts. Alternatively, after RCA the loop region can be hybridized with a sequencing primer and *in-situ* sequenced using repurposed Illumina sequencing reagents (13). Each sequencing round involves the incorporation of reversibly terminated fluorescently labeled nucleotides, washing, imaging and dye cleavage to allow the incorporation of the next nucleotide (**Fig. 4b**). With 4-colour chemistry, up to 4*^n^* transcripts can be multiplexed, where *n* is the number of sequencing rounds (13). Image processing is required to extract the location and identity of individual transcripts via image registration, RCP (spot) detection, and barcode decoding (**Fig. 4b**).

**Figure 4.**
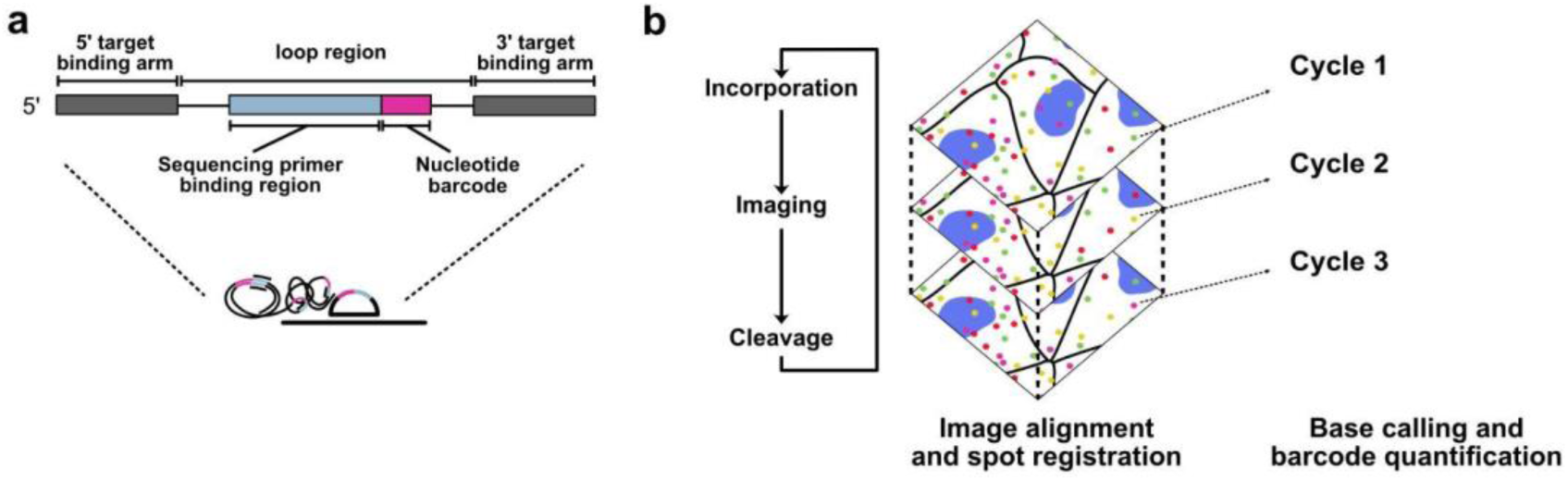
mudRapp Seq workflow for in situ RNA detection and quantification. (a) The loop region of the padlock probe can be customized with target-specific nucleotide barcode and by hybridizing sequencing primers to the rolling circle products (RCPs) and employing Illumina 4-colour chemistry for in situ sequencing. (b) These barcodes can be detected and quantified via sequencing. After sequencing primer hybridization, incorporation mix is added and the sample is imaged, followed by cleavage of the dye attached to nucleotide to prepare the sample for the next round of incorporation. After multiple sequencing cycles, maximum intensity projection is performed on z-stack images, followed by image alignment and spot registration. The fluorescent spots are then basecalled for each sequencing cycle to determine the barcodes. Finally, the spatial localization of detected targets is registered, enabling further quantifications.

To establish four colour *in situ* sequencing, we first examined the spectral properties and crosstalk of each nucleotide in their respective emission channels. We prepared four samples, each incorporating a different nucleotide on the first round of sequencing (**Supp. Fig. 5a**). The four dyes were well discriminated, with only minor bleed-through of the A nucleotide emission to the emission filter selected for T, which we corrected using a bleed-through filter during analysis (**Supplementary Fig. 5a**).

We next evaluated the cleavage and incorporation efficiency over six rounds of sequencing on well-defined barcodes (**Supplementary Fig. 6a**). By following RCP spots correctly base-called in the first round of sequencing, we found that nearly 99% of spots were correctly called in the next cycle of sequencing (99.3%, 99.3%, 98.8%, 97.6%, and 98.0% for cycles 2, 3, 4, 5 and 6 respectively) (**Supplementary Fig. 6b**). This analysis assumes each spot has only one signal per sequencing round (A,C,G,T). However, when distinctly barcoded RNA molecules are within the diffraction limit, it is possible for multiple channels to contain a signal within a single sequencing round. This can occur when RNA molecules are crowded in cells, but may also occur during the late stages of influenza A virus assembly when vRNAs associate together (2).

To address this, we tested a stringent decoding strategy that allows for multiple channels to be active (or completely absent) per sequencing round. In this control setting, we decoded greater than 92% of spots after 2 rounds of sequencing, and 84% spots after 6 rounds of sequencing defining the background of our assay (**Supplementary Fig. 6c**). We classified the ambiguous spots into ‘missing’ (spot missing the expected signal in any round of incorporation), ‘additional’ (the spot is assigned to another nucleotide in any round of incorporation), and ‘missing and additional’ (there is at least one round of sequencing where the expected nucleotide is missing according to the barcode and at least one round of sequencing where an unexpected nucleotide was called) (**Supplementary Fig. 6c**). Incorrect decoding was almost exclusively ‘additional’, likely due to the accumulation of signals from incomplete reactions during sequencing. This issue can be mitigated by restricting the number of sequencing cycles, or by using error-correcting barcodes.

### Temporal dynamics of Influenza A virus mRNAs and vRNAs

Having established mudRapp-seq, we applied this technique to study the replication dynamics of IAV vRNAs and mRNAs during an infection time course. Three RNA species are produced in a tightly controlled manner: viral RNA (vRNA), viral mRNA (mRNA), and complementary RNA (cRNA) (29–32). mRNA is produced early in infection until sufficient proteins are produced to allow replication of the vRNA, which occurs through a positive sense cRNA replication intermediate (32,33).

To allow the simultaneous differentiation of all eight vRNA and viral mRNA segments we engineered PLPs to include a two-nucleotide barcode (**Fig. 5a** and **Supplementary Data 4**). The first round of sequencing distinguishes between the vRNA and mRNA whereas the second round is used to identify specific individual segments (**Fig. 5a**). We infected MDCK cells with the PR8 strain at MOI of 0.3 and 1.0, and prepared samples from 0-8 h with an interval of 1 h (**Fig. 5b**). For the 1 MOI infection, mRNA expression could be detected from 1 h post-infection onward and it reached steady state at approximately 3-4 h (**Fig. 5c**). mRNAs exhibited a monophasic expression profile and were predominantly localized to the cytoplasm (**Fig. 5c**). In contrast, vRNA abundance was bi-phasic, with an initial increase seen at 1 h post-infection, and a second increase at 7 h post-infection (**Fig. 5c**). The initial phase was restricted to the nucleus whereas the second phase was largely driven by the appearance of vRNA in the cytoplasm (**Fig. 5c**). For 0.3 MOI of infection, a similar picture emerges except that the initial accumulation of mRNA was slightly delayed, and a second mRNA wave was seen at 7 h post-infection, likely driven by the infection of previously uninfected cells, although this remains to be tested (**Fig. 5c**). We further note that padlock probes cannot differentiate between mRNA and cRNA molecules as they bind to internal sequences common to both. However, we do not believe this affects these conclusions since the amount of cRNA is very low compared to mRNA, and cRNAs are localized predominantly in the nucleus, whereas mRNAs are present in the cytoplasm.

**Figure 5.**
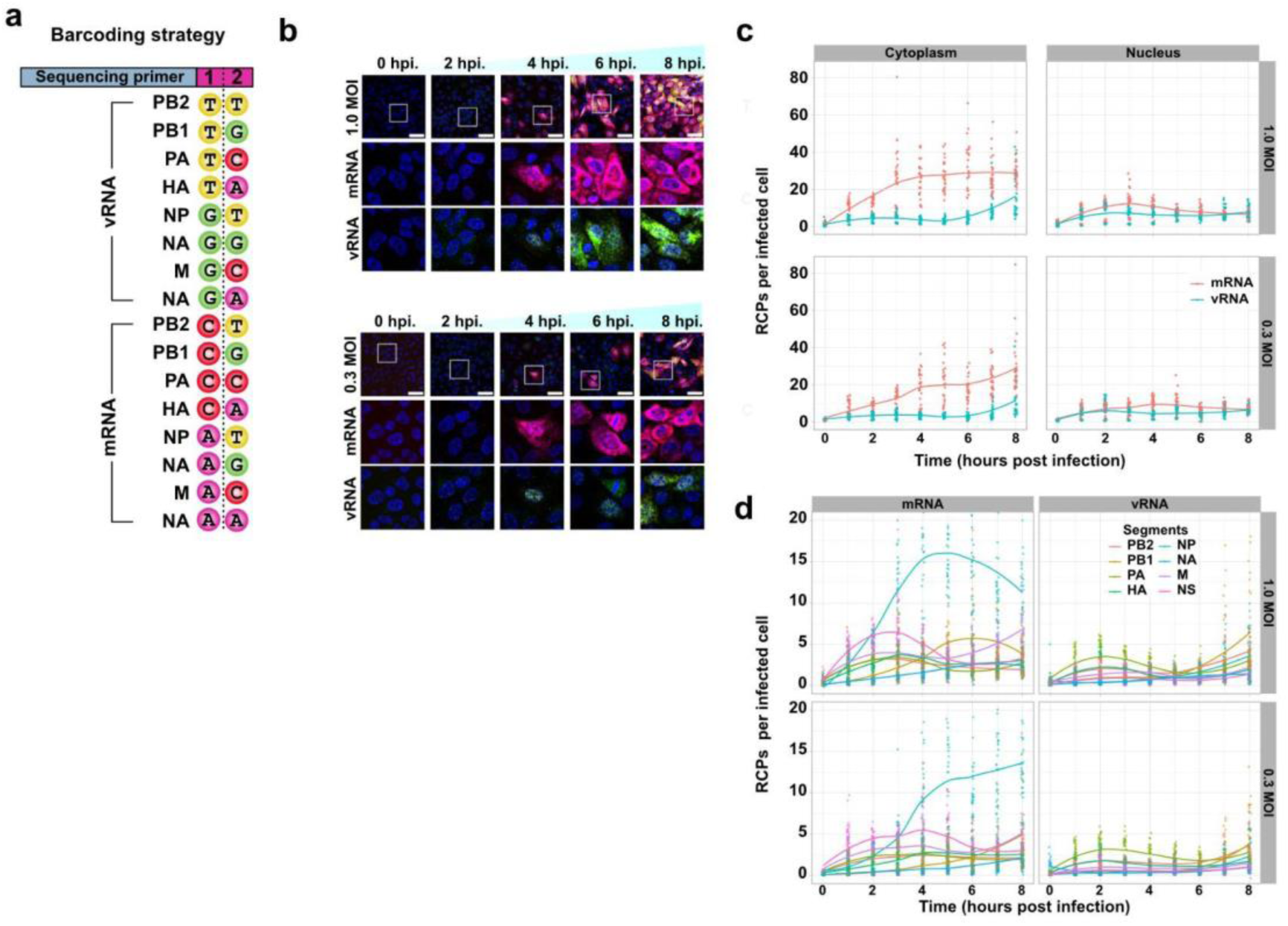
Simultaneous detection of the vRNA and viral mRNA in infected cells. (a) Barcode design on the padlock probe for simultaneous detection of the vRNA and mRNA in cells. The first round of incorporation distinguishes between the vRNA and mRNA molecules in infected cells and in combination with the second round of incorporation specific segments can be identified. (b) Representative image of PR8 infected MDCK-cells with 0.3 MOI and 1 MOI after the first round of incorporation where the vRNA and mRNA can be distinguished. (Scale bar = 50 µm). (c) Graph showing the bulk transcription and replication pattern of PR8 infected with 0.3 and 1 MOI for 0-8 hpi Points are individual (field of view) FOVs, lines are locally estimated scatterplot smoothing (loess) smoothedconditional means. (d) Distribution of the single vRNA and mRNA segments at 0-8 h for 0.3 and 1 MOI infected cells. Points are individual FOVs, lines are loess smoothed conditional means. y-Axis is limited to range 0-20, some points exceed this range and are therefore not shown, all points are included in calculating the smoothing.

Analysis of individual segment mRNA expression revealed significant segment-specific variations (**Fig. 5d** and **Supplementary Fig. 7**). NP mRNA was the most abundant mRNA, presumably because NP is required in higher quantity for vRNP formation (**Fig. 5d** and **Supplementary Fig. 7**). Notably, one of the earliest mRNAs expressed was NS, which is alternatively spliced into NS1 and NS2/NEP (34) (**Fig. 5d** and **Supplementary Fig. 7**). NS1 is the predominant product, which acts as the primary viral antagonist for the host cellular responses and is likely to be required during early stages of infections (34). mRNAs encoding the IAV structural proteins required for virion formation and budding, such as M (alternatively spliced to form the M1 structural protein and the M2 transmembrane proton channel), HA (hemagglutinin), and NA (neuraminidase) were increasingly transcribed towards the later stages of the infection cycle (from 4-h post-infection) (**Fig. 5d** and **Supplementary Fig. 7**). Interestingly, the expression of M mRNA was highly correlated with the time series for total vRNA at both MOIs (MOI 1, Pearson’s r=0.96, p-value <0.0001; MOI 0.3, Pearson’s r=0.89, p-value = 0.001), which could indicate an involvement of M1 or M2 protein expression with the switch from mRNA transcription to vRNA replication (**Supplementary Fig. 8** and **Supplementary Data 4**).

Most RCPs could be assigned to a single mRNA or vRNA segment. Intriguingly, however, the number of ambiguous RCPs rose steadily from 4 and 6 hpi, for 0.3 and 1 MOI, respectively (**Supplementary Fig. 9a**). Ambiguous spots were detected mainly in the cytoplasm, which could be due to the presence of vRNA complexes during their transport to assembly sites at the plasma membrane (**Supplementary Fig. 9a**). To investigate this further, we used the first sequencing round to classify ambiguous RCPs into mRNA only, vRNA only or mixed (containing both vRNA and mRNA). All three ambiguous species monotonically increased at each time point for the 0.3 MOI infection, as expected of molecular crowding. However, at late time points in the 1.0 MOI case, the number of ambiguous vRNA spots increased, while, contrary to expectation from crowding, the number of mRNA-multi spots decreased (**Supplementary Fig. 9b**).

In sum, our data supports a model where incoming vRNPs are first transcribed into mRNAs in a non-stoichiometric manner. Only later are vRNAs replicated, and the onset of replication correlates with the expression of M mRNAs, suggesting their role in regulating the switch from transcription to replication.

### Segment-specific associations with infection heterogeneity

Infection outcomes depend on the physiological states of the different cells within an organism (35–41), which cannot be explored using bulk measurements of viral RNA. We therefore established a single-cell analysis pipeline using semi-supervised machine learning to localize RCPs to individual infected cells (**Fig. 6a**).

**Figure 6.**
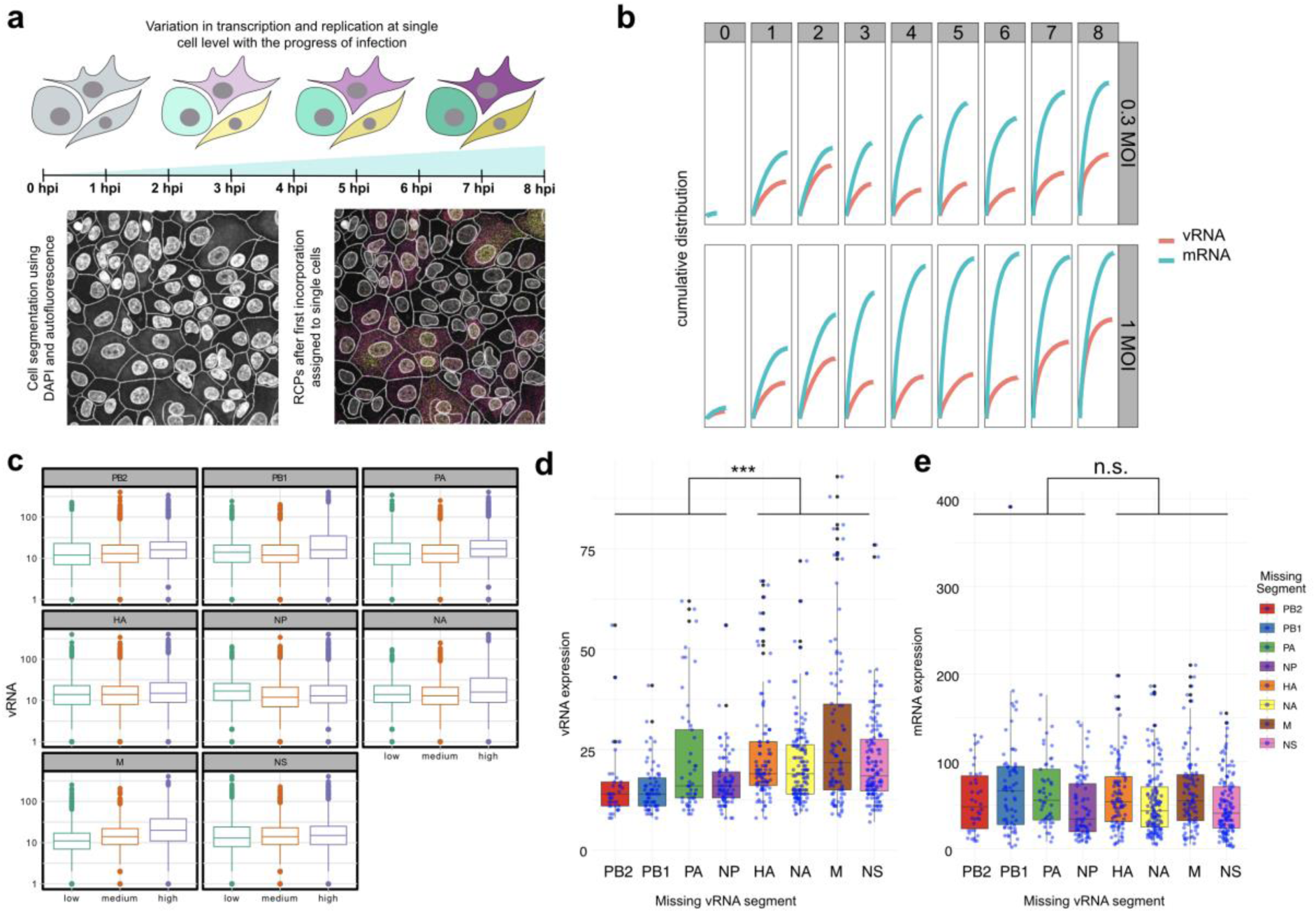
Single-cell analysis of transcription and replication heterogeneity during early infection. (a) Diagram of variations in transcription and replication at single-cell level at early time points of infection. We fixed the cells every hour till 8 h post-infection to capture the variation in transcription and replication every hour at the single-cell level. For single-cell visualization of the RCPs after in-situ sequencing, cell and nucleus segmentation is performed. (b) Cumulative distribution of total vRNA and mRNA at each time point of infection. The number of cells included is on the x-axis and the total number of RCP spots in these cells on the y-axis, cells are sorted descending by number of RCPs. (c) Boxplot illustrating the levels of total vRNA on a log-scale in cells categorized as low, medium, or high expressers for individual mRNA segments. (d-e) Abundance of total vRNA (d) and mRNA (e) in cells at hpi>5 missing exactly 1 vRNA segment. x-axis shows the missing segment, and the y-axis the total sum of vRNA and mRNA in cells. Statistical test was Wilcoxon rank sum test with continuity correction.

We first performed a cumulative sum analysis of the total number of mRNA or vRNA molecules detected per cell across different MOIs and time-points (**Fig. 6b**). This analysis revealed significant cell-to-cell heterogeneity in vRNA and mRNA levels, which was also observed for individual segments (**Supplementary Fig. 10**). Such variability suggests that viral transcription and replication occur unevenly across infected cells. To quantify this heterogeneity, we calculated Gini coefficients. Gini’s coefficient (or Gini index) is a statistical measure of inequality or dispersion in a dataset. It ranges from 0 to 1 where, 0 stands for perfect equality (all cells contain the same number of transcripts) and 1 stands for perfect inequality (all transcripts are concentrated in a single cell) (42). Both mRNA and vRNA showed increasing Gini coefficients over time, with the largest increases occurring at 3-4 hpi for mRNA and 6-7 hpi for vRNA. These peaks coincided with increased abundance observed in the bulk analysis. These findings suggest that as viral transcription and replication intensify, certain cells accumulate disproportionately high transcript levels, potentially due to differences in viral entry, replication efficiency, or host cell factors (**Supplementary Fig. 11**).

We next investigated whether heterogeneity in overall vRNA abundance could be related to the specific expression of a particular segment. Focusing on cells expressing all eight mRNAs at late time points (>5 hpi), we used a cumulative sum analysis to categorize cells into low, medium, and high mRNA expressing groups for each segment. Examining the relationship between these categories and vRNA abundance suggested that high mRNA expressing cells generally expressed higher amounts of vRNA (**Fig. 6c**). To test the significance of these associations, we developed a linear model that uses mRNA as inputs to predict the total vRNA (**Table 2**). This analysis identified significant positive associations between mRNA expression of PB1, PA, NA, and M with vRNA abundance, with the most significant association for M mRNA (p<0.0001), again suggesting a potential role for M vRNP segment proteins in regulating the switch from transcription to replication.

**Table 2:** Linear model using mRNAs as inputs to predict the total vRNA. These values have been filtered for cells with at least 40 mRNA spots of at least 6 distinct mRNAs. Based on this PB1, NA, NP, M, and PA contribute statistically significantly (the contribution of NP is small and negative).

|  | Term | Estimate | Std. error | Statistic | p. value |
| --- | --- | --- | --- | --- | --- |
| 1 | intercept | 2.17 | 0.913 | 2.38 | 1.73e- 2 |
| 2 | mRNA_PB2 | -0.203 | 0.132 | -1.54 | 1.24e- 1 |
| 3 | mRNA_PB1 | 0.500 | 0.0562 | 8.89 | 9.87e-19 |
| 4 | mRNA_PA | 1.16 | 0.0884 | 13.1 | 1.68e-38 |
| 5 | mRNA_HA | -0.0872 | 0.0740 | -1.18 | 2.39e- 1 |
| 6 | mRNA_NP | -0.142 | 0.0229 | -6.21 | 5.88e-10 |
| 7 | mRNA_NA | 1.34 | 0.129 | 10.4 | 6.75e-25 |
| 8 | mRNA_M | 0.943 | 0.0462 | 20.4 | 3.15e-87 |
| 9 | mRNA_NS | 0.0479 | 0.0641 | 0.748 | 4.55e- 1 |

To further investigate this, we examined the expression of M1 and M2 proteins through western blotting and immunofluorescence imaging. At the MOIs used in this study (0.3 and 1), minimal M1 protein expression was observed during the early stages of infection, consistent with the low levels of M mRNA under these conditions. At a higher MOI of 5, western blot analysis detected M1 protein as early as 5 hpi, whereas M2 was detected only at 7 hpi (**Supplementary Fig. 15a**). Immunofluorescence staining showed a similar pattern (**Supplementary Fig. 13 and 14**), confirming that M1 is expressed earlier than M2. To further explore the role of M1 and M2 in viral transcription and replication, we performed RT-qPCR to quantify vRNA and vmRNA in cells pre-expressing RNP alone or in combination with M, M1, or M2. RNP pre-expression increased both vRNA and mRNA abundance, which was further enhanced by co-expression of M (**Supplementary Fig. 15b**). Notably, RNP+M1 pre-expression resulted in the highest levels of both vRNA and vmRNA, whereas RNP+M2 pre-expression led to a strong reduction in vRNA and vmRNA levels (**Supplementary Fig. 15b**). Thus, M1 and M2 proteins have different effects on vRNP and vmRNA: M1 enhances both transcription and replication whereas M2 negatively altered vRNA and vmRNA synthesis.

While the above analysis focused on cells where all eight segments were detectable, many cells contained an incomplete set of vRNAs, especially when infections were carried out at lower MOI (**Supplementary Fig. 12**). To explore how missing vRNAs impact viral replication, we filtered for cells expressing 7 vRNA segments at late time points (> 5 hpi). We found that lack of PA, PB1, PB2, and NP vRNAs were strongly associated with lower overall vRNA abundance, as expected from their involvement in vRNA replication (**Fig. 6d**). Interestingly, we did not find a strong statistical association between the levels of PB1, PB2, PA, and NP vRNAs and total mRNA levels. This suggests that the presence of NP and polymerase-encoding vRNAs is not strictly required for mRNA transcription, likely because transcription can occur from polymerase complexes already present on incoming vRNPs during infection (**Fig. 6e**). To further investigate this, we analyzed cells that expressed only seven mRNA segments (rather than vRNAs), allowing us to more directly assess the role of newly synthesized NP and polymerase in mRNA production. We found that the absence of PB1, PB2, PA, or NP mRNA did not significant correlate with lower total mRNA levels, again suggesting transcription from incoming vRNPs can be efficient in the absence of newly synthesized polymerase proteins (**Sup. Fig. 11**) (43). However, we do not exclude the possibility that a secondary transcription phase occurs following vRNP replication, which we may not have had the statistical power to detect.

## Discussion

The dynamics of viral infections at the cellular level have long been obscured by the limitations of bulk analysis techniques. While these techniques have provided valuable insights into average viral behaviors across populations of cells, bulk analyses mask significant cell-to-cell variability that is increasingly recognized as a critical factor in infection outcomes.

The most widespread approach addressing this challenge is single-cell RNA sequencing (scRNA-seq). This is based on including cell-specific unique molecular identifiers (UMI) to cDNA molecules during reverse transcription of polyadenylated RNAs (44,45). This has allowed the analysis of the viral and host response during IAV infection, revealing heterogeneity in mRNA production (41) and the effect of viral factors on the host response (39,46,47). However, the use of UMI-oligo(dT) primers in scRNA-seq assays has limited the view of viral behavior to the viral mRNAs, as vRNAs do not have poly-A tails, leaving the co-dynamic expression of vRNA and mRNA poorly understood at the single-cell level.

Here, we established mudRapp-seq to simultaneously visualize the transcription and replication of all eight IAV segments. As a spatial transcriptomics method, mudRapp-seq allows counting RNA molecules in cells, whilst maintaining information about their localization. Our approach borrows strengths from different spatial transcriptomics techniques whilst avoiding many of their key limitations. Similar to smFISH, mudRapp-seq employs multiple probes per target to boost detection efficiency. However, we achieve signal saturation at approximately 5-6 PLPs per RNA target, compared to the 20-50 unique probe binding sites per RNA required by smFISH. This is due to signal amplification achieved through RCA of successfully bound and ligated PLPs. This feature makes mudRapp-seq particularly suited for short RNAs, where sufficient smFISH probe targets are not available. Another advantage of mudRapp-seq is its ability to co-image genetically similar molecules, such as closely related circulating IAV viruses, thanks to the target specificity provided by the PLP ligation step. This level of discrimination is not achievable with smFISH to the lack of unique probe binding sites. While PLP-based methodologies using in-situ reverse transcription typically balance high specificity with poor sensitivity, direct RNA binding PLPs improve efficiencies by bypassing that reverse transcription step. Additionally, multiple PLPs help mitigate probe binding issues caused by RNA structure within the target. As chemical probing clearly demonstrated the inhibitory role of RNA structure, future optimizations should account for target structure in probe design, as well as the ligation junction, enzyme, buffer, and hybridization arm size, which may allow less probes per target to be used to reduce cost without sacrificing efficiency (6,16).

Using mudRapp combined with in situ sequencing, we followed all eight vRNA and mRNA of IAV from just after entry to replication in cells. From these measurements, we observed that viral mRNA expression starts as early as 1 h post-infection, whereas vRNA replication initiated at 7 h post-infection, with different kinetics for each segment. This difference in transcription and replication timing suggests a coordinated regulation of viral RNA processes, orchestrated by viral and host factors (33,48–52). Furthermore, our analysis indicated segment-specific regulation of mRNA transcription, indicating a control mechanism governing viral mRNA expression, perhaps regulated by secondary structures in the mRNA that alter the transcription and/or stability of the individual mRNA (53). In contrast to mRNA, vRNA replication proceeded relatively uniformly across all the viral RNA segments.

Strikingly, we identified a temporal correlation between the onset of vRNA replication and expression of mRNA from the M segment. M mRNA codes for two proteins: M1 and M2, with M2 being expressed from an alternatively spliced form of the M mRNA (54). Both proteins have pleiotropic functions and are essential for viral replication and virus assembly, but the M2 protein expression matched most closely the switch to vRNA replication at the beginning at 7 hpi. Moreover, pre-expression of RNP+M1 led to higher production of vRNA and vmRNA upon infection compared to WT, whereas pre-expression of RNP+M2 resulted in a strong decrease in vRNP and vmRNA abundance. We believe that these data better support a role for M2 in switching off transcription, although further follow up studies will be required to define the role of M proteins in polymerase activity.

Finally, we performed a single-cell analysis of mRNA and vRNA production, finding extensive cell-to-cell heterogeneity. As has been described by others, we observed a substantial proportion of cells that failed to replicate all vRNA segments (41,55). Incomplete expression was less frequent at high MOI, in agreement with the idea that higher viral loads may enhance the efficiency of vRNA replication through complementation (24,56,57). Interestingly, we found that cells missing either component of the polymerase complex vRNA segments or NP were associated with almost negligible replication of the vRNA. However, these same missing segments had less significant effect on levels of mRNA. The observed association between missing polymerase segments and NP segment does not necessarily imply a causal relationship. However, one plausible explanation could be that incoming vRNPs, which are already loaded with the polymerase complex, are programmed for transcription, whereas newly synthesized polymerases and NP proteins are required for vRNA replication. Along these lines, it is reported that inhibition of translation with cyclohexamide prevents vRNA replication without affecting mRNA transcription, but this defect is rescued when PB1, PB2, PA, NP are pre-expressed in cells (43). To further explore the role of polymerase, NP and M in the replication cycle, it would be useful to be able to selectively degrade specific newly synthesized IAV proteins during a replication cycle.

In summary, our newly established methodology offers a powerful tool for dissecting the molecular mechanisms underlying IAV virus replication and pathogenesis. Future applications of this technique may elucidate the expression of various host factors, such as interferon-stimulated genes, in infected cells and their neighbors. Additionally, this approach could shed light on subcomplex formation and packaging pathways, further enhancing our understanding of viral replication strategies. By providing a high-resolution view of viral RNA dynamics, our work contributes to a more comprehensive understanding of influenza A virus biology and contributes to our understanding of viral pathogenesis and evolution. Moreover, enhanced multiplexed RNA detection within individual cells will more generally enhance our understanding of transcriptional heterogeneity and the spatial localization of gene expression changes, potentially providing insights into other diseases such as cancer and neurodegenerative disorders.

## Acknowledgments

We would like to thank Simone Backes and Anke Sparmann for critical feedback. We also thank Liqing Ye, Marco Olguin-Nava, Charlene Börtlein, and Patrick Bohn for meaningful discussion and technical assistance.

## Author contributions

R.P.S. and S.A. conceived the study. S.A. performed the imaging experiments. J.L. did the qPCRs, N.G., and S.M. performed the western blot and immunofluorescence imaging. S.A and A.S.G.B. performed the DMS probing and nanopore sequencing. M.J.A., J.S, R.P.S., S.C.F., and S.A. performed the analysis. U.B.A. helped with virus propagation. R.P.S., S.A and M.J.A. wrote the manuscript with contributions from the other authors.

## Funding

We thank the Helmholtz Association (VH-NG-1347 to RPS). RPS also acknowledges the interdisciplinary Thematic Institute IMCBio+, as part of the ITI 2021-2028 program of the University of Strasbourg, CNRS and Inserm, IdEx Unistra (ANR-10-IDEX-0002), SFRI-STRAT’US (ANR 20-SFRI-0012), and EUR IMCBio (ANR-17-EURE-0023) under the framework of the French Investments of the France 2030 Program. The funders had no role in study design, data collection, analysis, decision to publish, or preparation of the manuscript.

## Data and code availability

Raw data is deposited in the BioImage Archive (will be archived). All analysis code is openly available at https://github.com/BioMeDS/mudRapp-seq. A snapshot of this repository at time of submission is additionally archived at Zenodo (will be deposited).

## Competing interests

The authors declare no competing interests.

## Notes

### Competing Interest Statement

The authors have declared no competing interest.

