## Supplementary Data for "Visualizing the transcription and replication of influenza A viral RNAs in cells by multiple direct RNA padlock probing and *in-situ* sequencing (mudRapp-seq)"

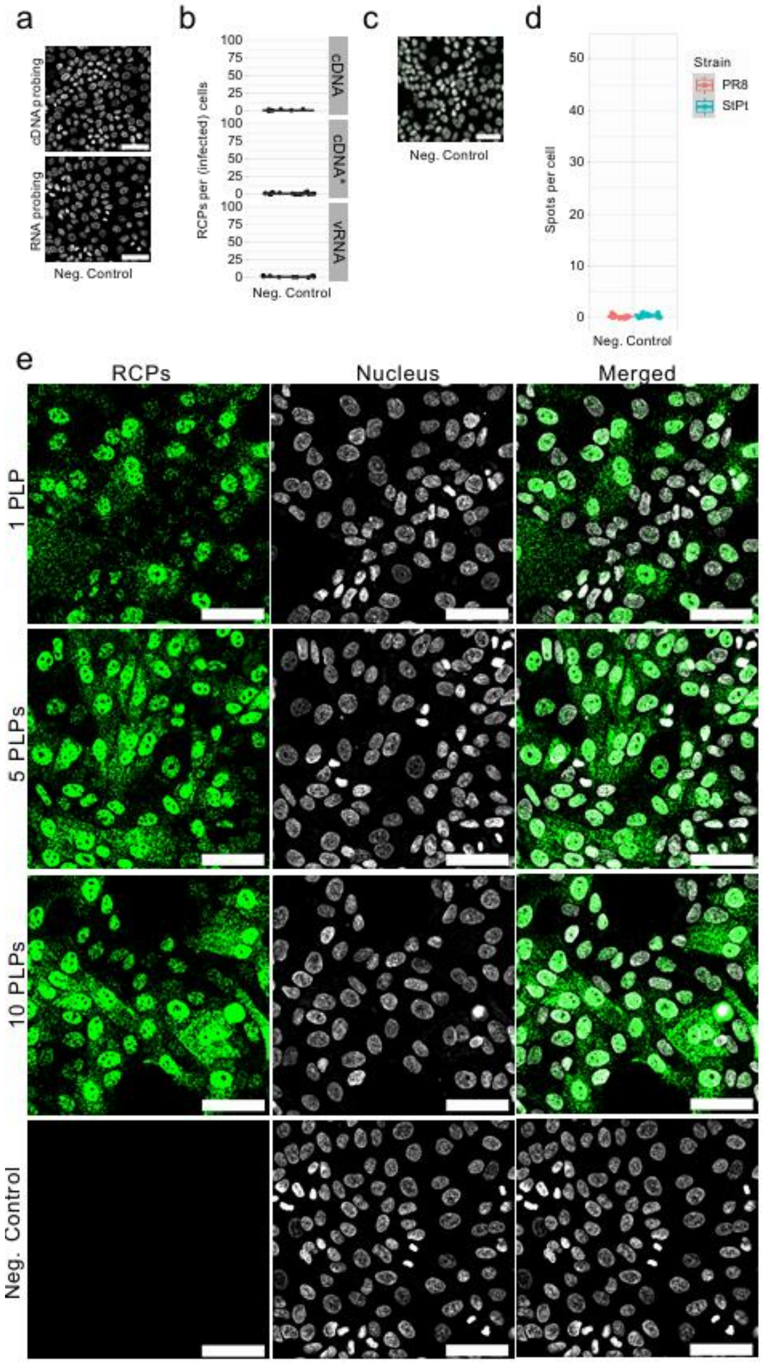


**Supplementary Figure 1**. **Specificity assessment of padlock probes (PLPs) in uninfected MDCK cells.** (a) Uninfected MDCK cells were probed with all 10 PLPs for both cDNA and direct RNA probing with negligible background (Scale bar = 50 µm). (b) Quantification demonstrated minimal unspecific binding of the PLPs. Range on the y-axis (0-100) matches that of figure 1c (c) Uninfected MDCK cells were probed with the PLP set for PR8 and St. Petersburg H1N1 strains (scale bar = 50 µm) (d) The quantification shows negligible nonspecific binding of padlock probes. Range on the y-axis (0-45) matches that of figure 1e. (e) The Zoomed images the direct RNA padlock probing images of figure 1b in individual imaging channels, namely for RCPs and DAPI for nucleus and the merged RCPs + Nucleus images.

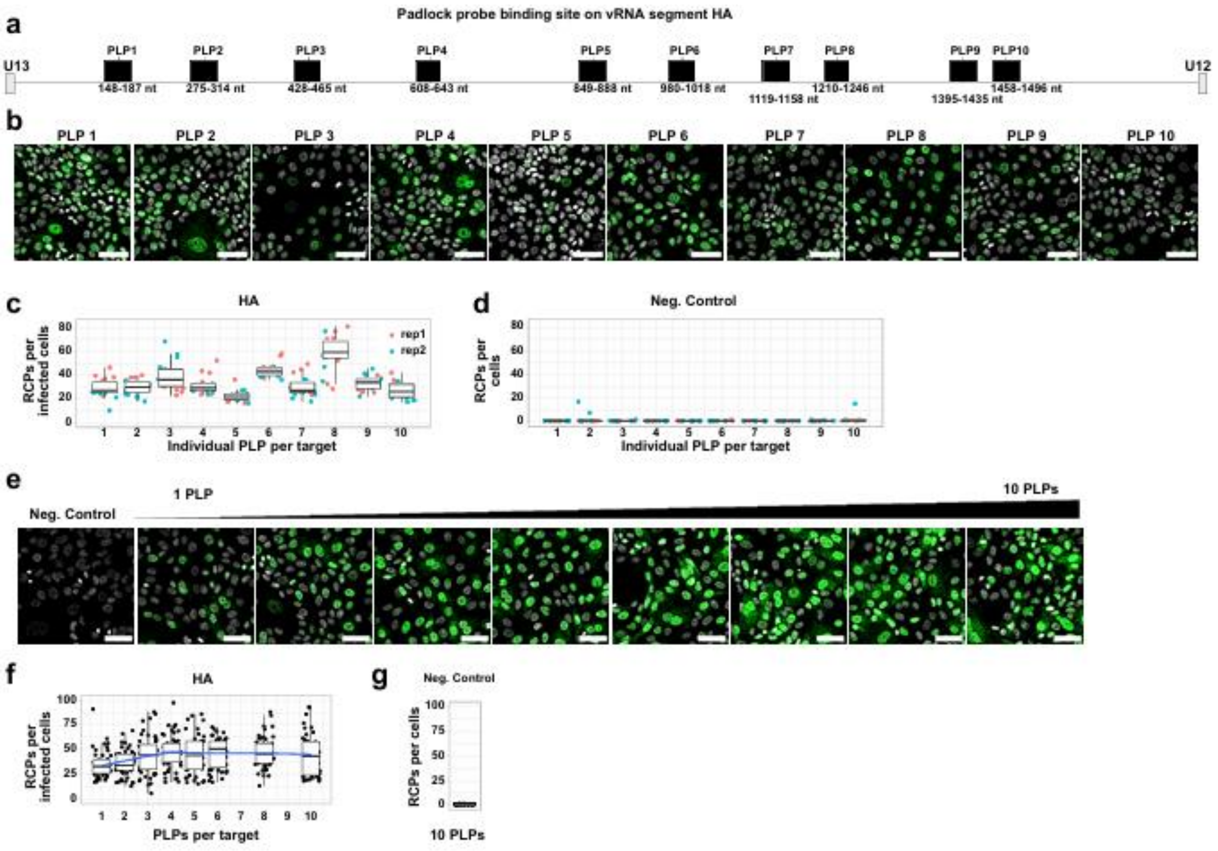


**Supplementary Figure 2**. **Multiple-direct RNA padlock probing improves the detection efficiency of the RNA targets.** (a) Schematic representation of padlock probe (PLP) binding sites on the HA segment of IAV strain PR8 vRNA. (b) Images demonstrating heterogeneity in PLP binding across different regions of the HA segment (scale bar = 50 µm). (c) Binding efficiency of individual PLPs for the HA segment, showing variation in probe performance. (d) Quantification of unspecific binding of the individual padlock probe. Negative control was performed on non-infected MDCK cells. (e) Images of PR8-infected MDCK cells (1 MOI, 6 h post-infection) probed with an increasing number of padlock probes. (f) Graph showing the increase in target detection efficiency as a function of the number of padlock probes per target. The blue line represents smoothed data obtained through local polynomial regression fitting. (g) Negative control with all 10 HA PLPs on uninfected cells.

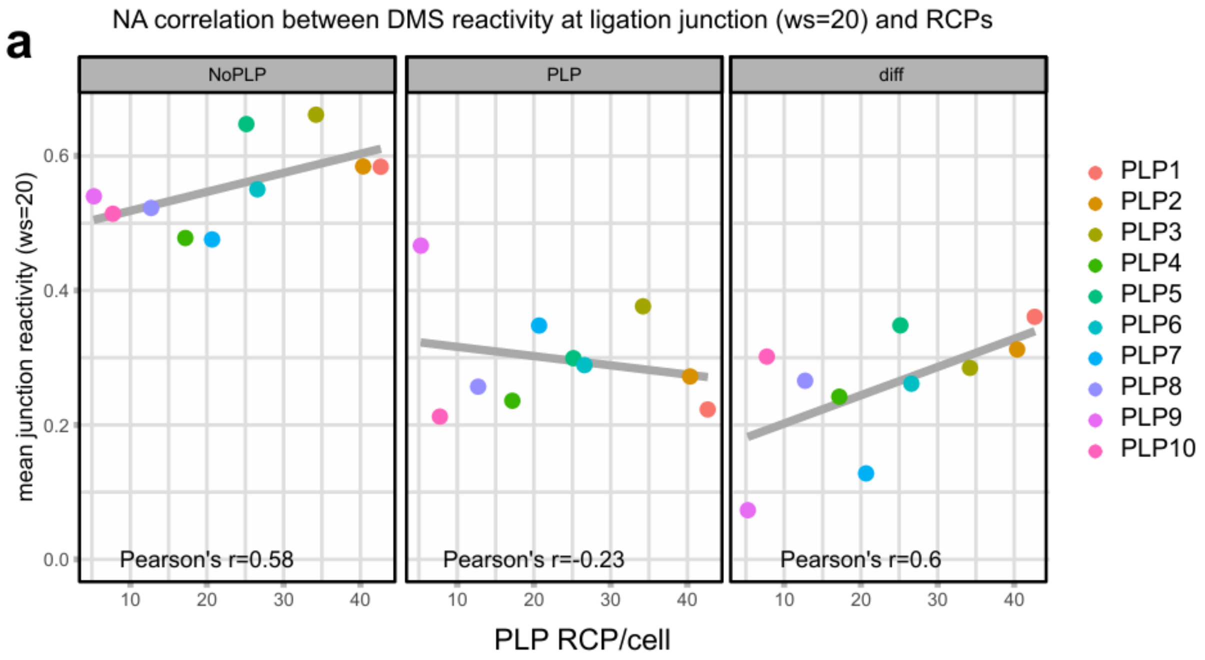


**Supplementary Figure 3. DMS reactivity correlation at the ligation junction of the PLPs on NA segment.** Correlation of binding efficiency (in RCP/cell) and DMS reactivity at target site with and without padlock probe binding and the difference between the two conditions at the ligation junction.

a

b

c

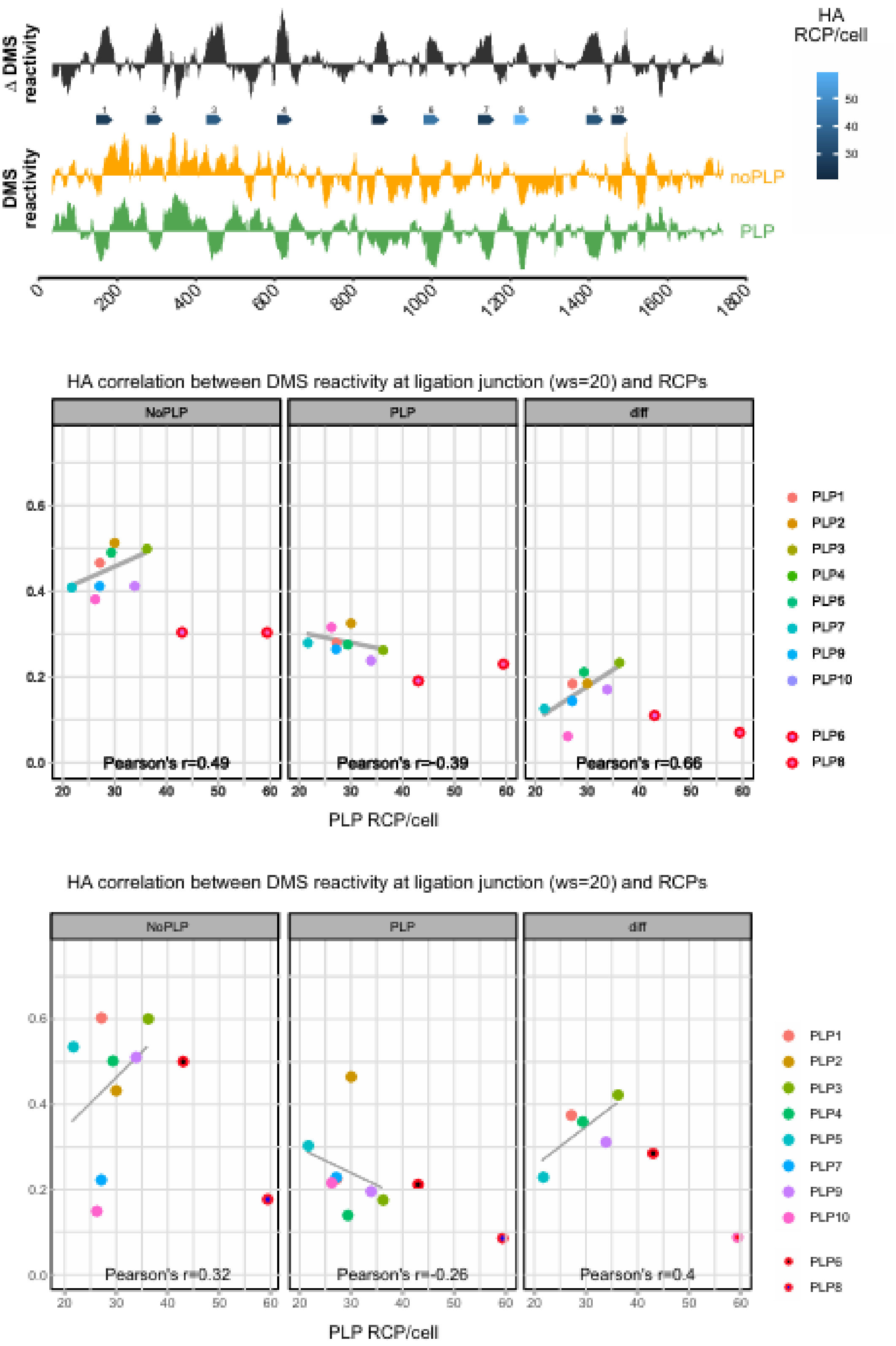


**Supplementary Figure 4. Role of RNA structure in the binding efficiency of the padlock probe on HA segment.** (a) DMS reactivity profile of the HA viral RNA segment with and without padlock probe binding. This graph illustrates how the presence of padlock probes affects the accessibility of RNA nucleotides to DMS modification along the length of the HA segment. Individual PLPs are displayed as blue arrows, colored by binding efficiency as measured by RCP/cell. Illustration generated with genomes. (b) Correlation analysis of DMS reactivity at target sites with and without padlock probe binding. The scatter plot shows the relationship between DMS reactivity in both conditions, with the difference highlighted to indicate structural changes upon probe binding. (c) Correlation of DMS reactivity at target site with and without padlock probe binding and the difference between the two conditions at the ligation junction of HA segment.

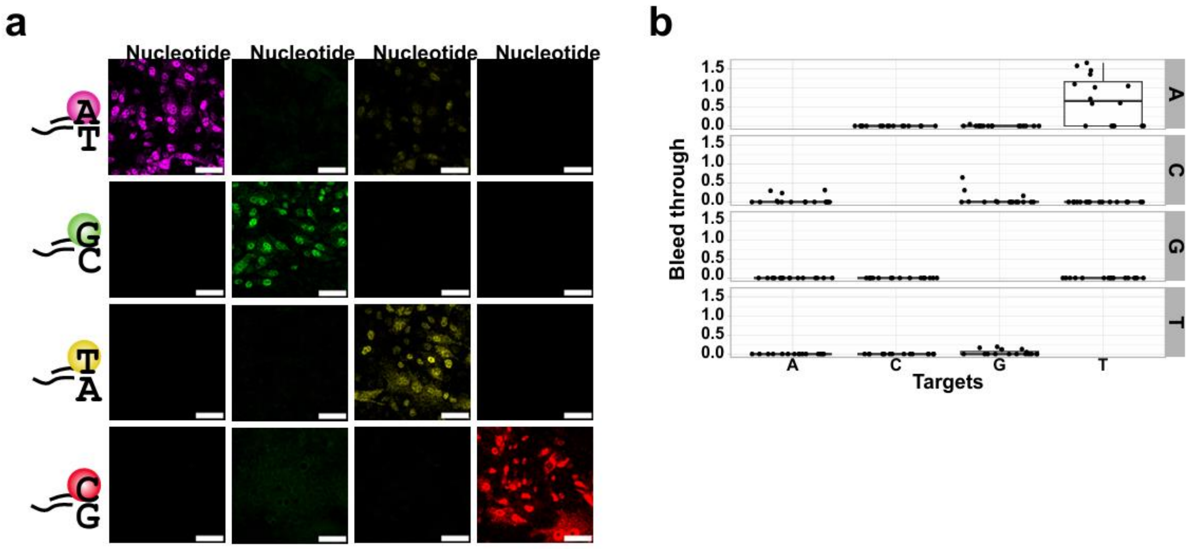


**Supplementary Figure 5. Nucleotide-specific fluorescence and channel crosstalk during *in-situ* sequencing.** (a) Representative images of four separate samples, each showing the result of the first round of nucleotide incorporation. In each sample, only one type of nucleotide (A, C, G, or T) was added. (Scale bar = 50 µm). (b) Quantification of fluorescence signal bleed-through between channels for each nucleotide as described by Maric et al.[1]. The plot demonstrates the specificity of each nucleotide-specific dye and the extent of crosstalk between channels. A minor bleed-through of the 'T' dye into the channel for 'A' nucleotide is observed.

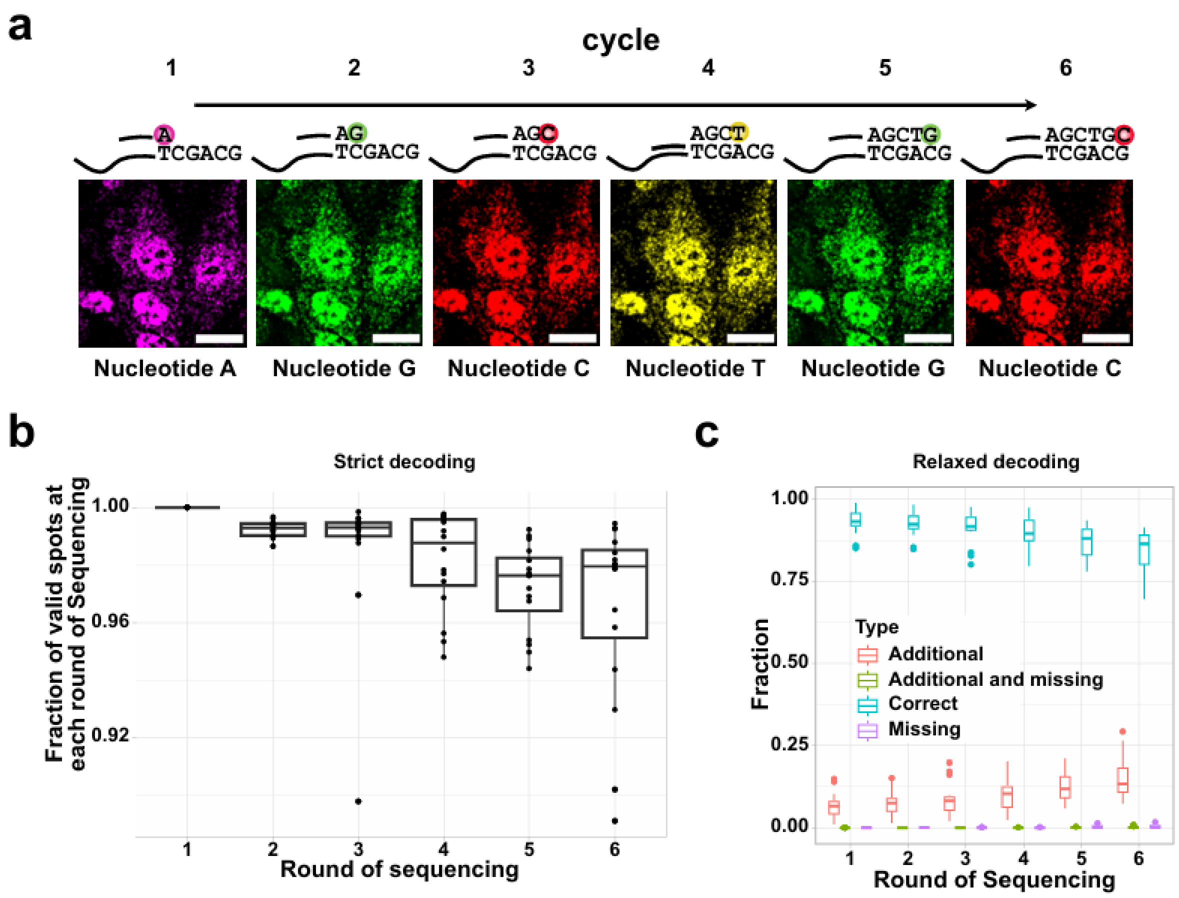


**Supplementary Figure 6. Sequencing accuracy and fidelity analysis of mudRapp-seq** (a) Example of images with six rounds of incorporation, where at each round only one nucleotide will be incorporated in the sample. (Scale bar = 50 µm) (b) The fraction of correct incorporations across six rounds of sequencing. Each percentage represents the proportion of spots that maintained correct base calling relative to the correct spots from the previous round, starting with the correctly called spots in round 1 (c) Quantification of correctly and incorrectly base-called spots (Rolling Circle Products, RCPs) after each round of incorporation, based on the expected barcode in the Padlock Probe (PLP). Incorrect spots include additional spots, missing spots, and combinations of additional and missing spots.

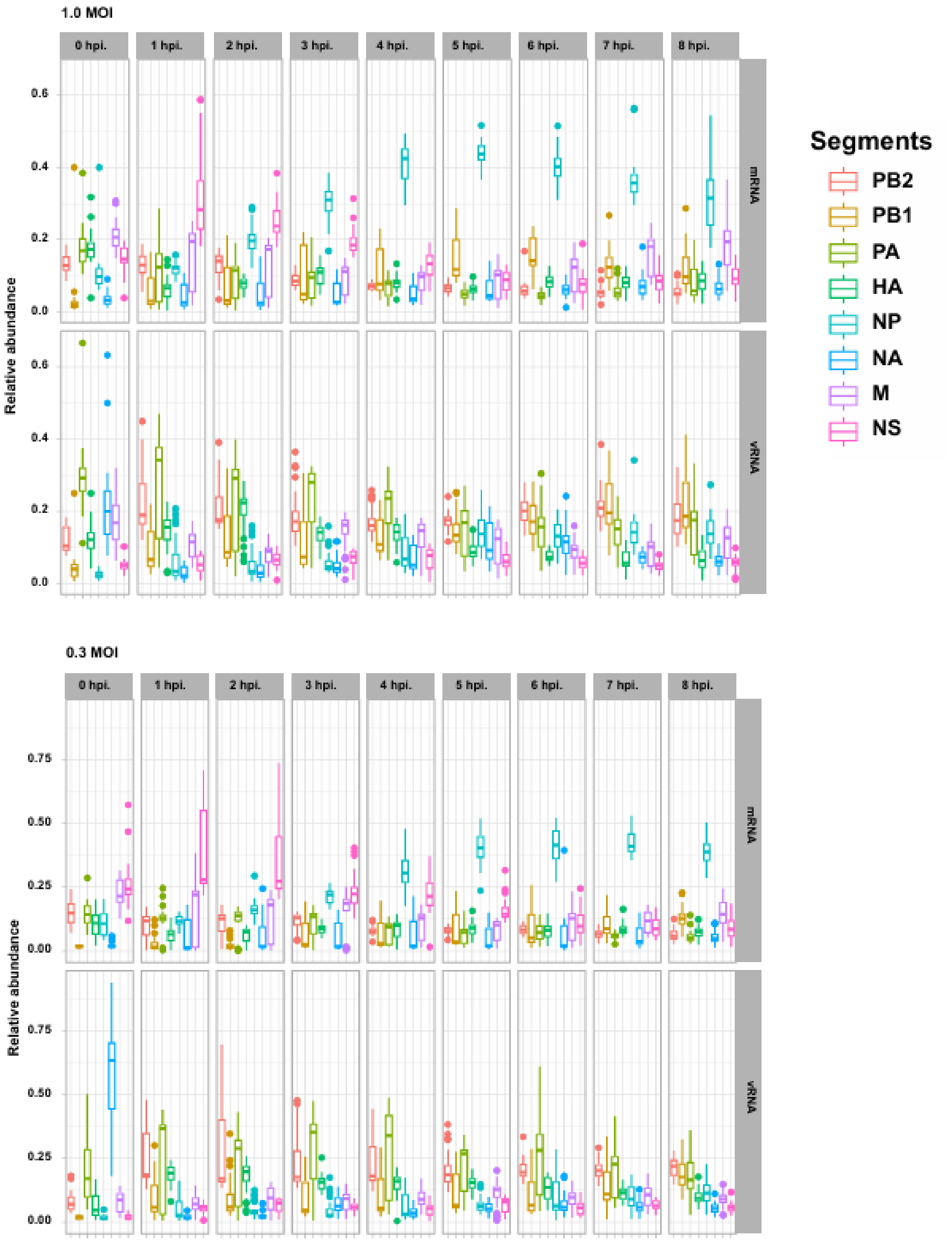


**Supplementary Figure 7. Temporal dynamics of Influenza A virus RNA segment abundance.** Relative abundance of all eight segments of both viral RNA (vRNA) and viral mRNA in infected cells across multiple time points post-infection. The number of spots that were detected was counted and normalized to the total number of spots per cell. This figure illustrates the temporal progression of viral genome replication (vRNA) and transcription (mRNA) for each of the eight influenza A virus segments for 0.3 and 1 MOI of infection.

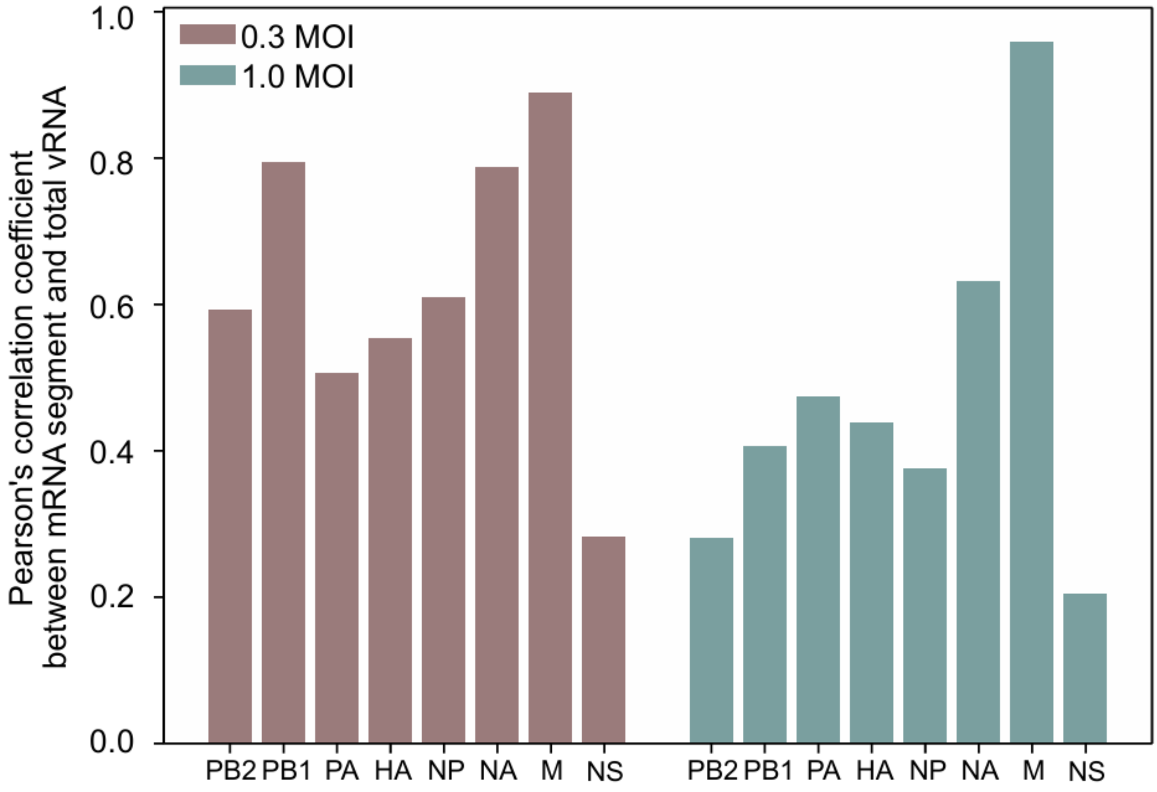


**Supplementary Figure 8. Correlation between M segment mRNA expression and total vRNA levels during Influenza A virus infection.** The plot shows Pearson's correlation coefficient for the time-course expression of each viral mRNA in relation to total viral RNA (vRNA) levels at 0.3 and 1 MOI of infection.

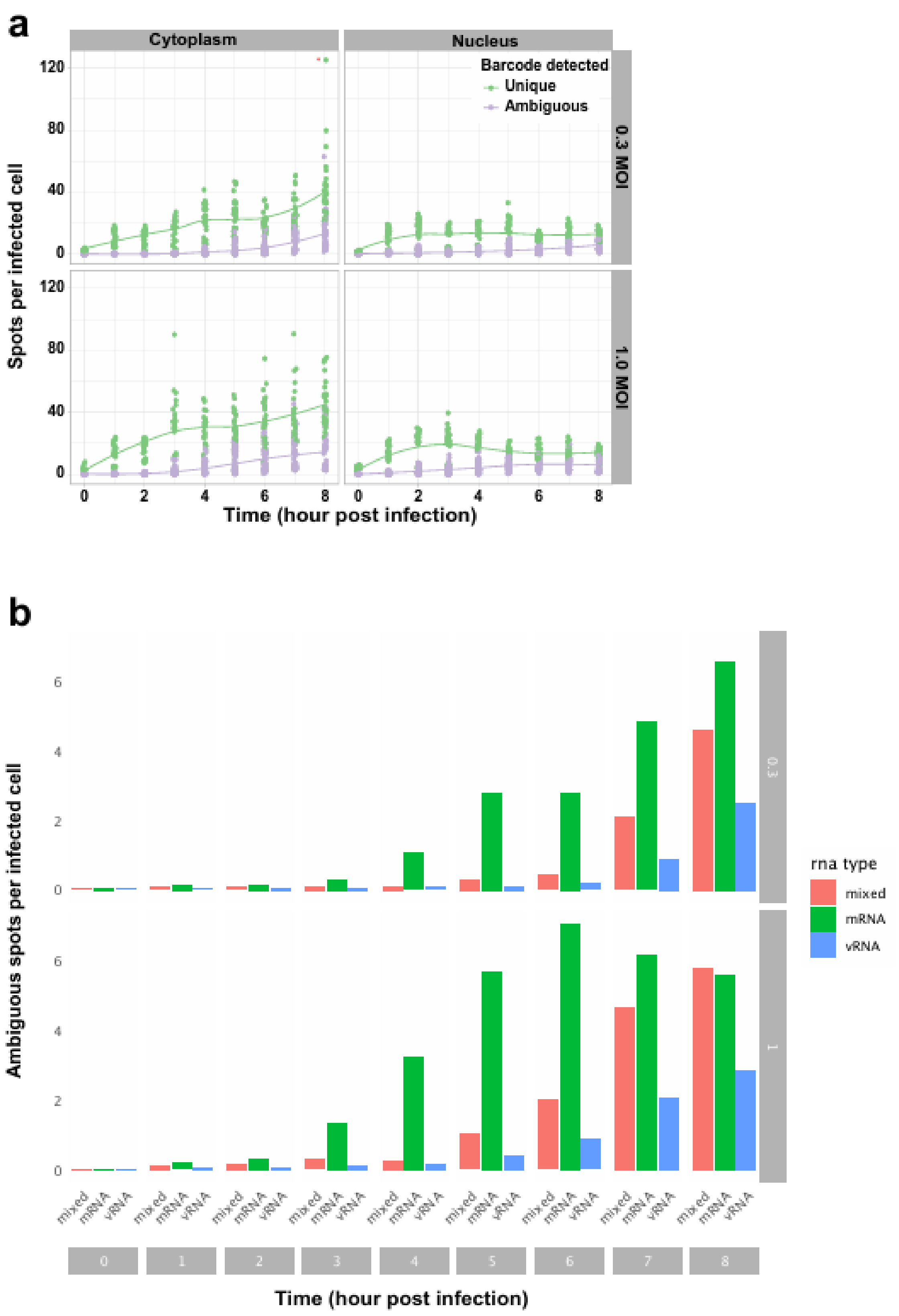


**Supplementary Figure 9. Unique and ambiguous Influenza A virus RNA species in infected cells** (a) This graph illustrates the temporal progression of viral RNA accumulation, distinguishing between uniquely identifiable RNA species and those with ambiguous classifications of total viral RNA and viral mRNA in infected cells from 0 to 8 h post-infection.

(b) Detailed breakdown of ambiguous RNA species in infected cells. This plot shows the relative proportions of ambiguous mRNA, vRNA, and mixed mRNA-vRNA populations across the infection time course.

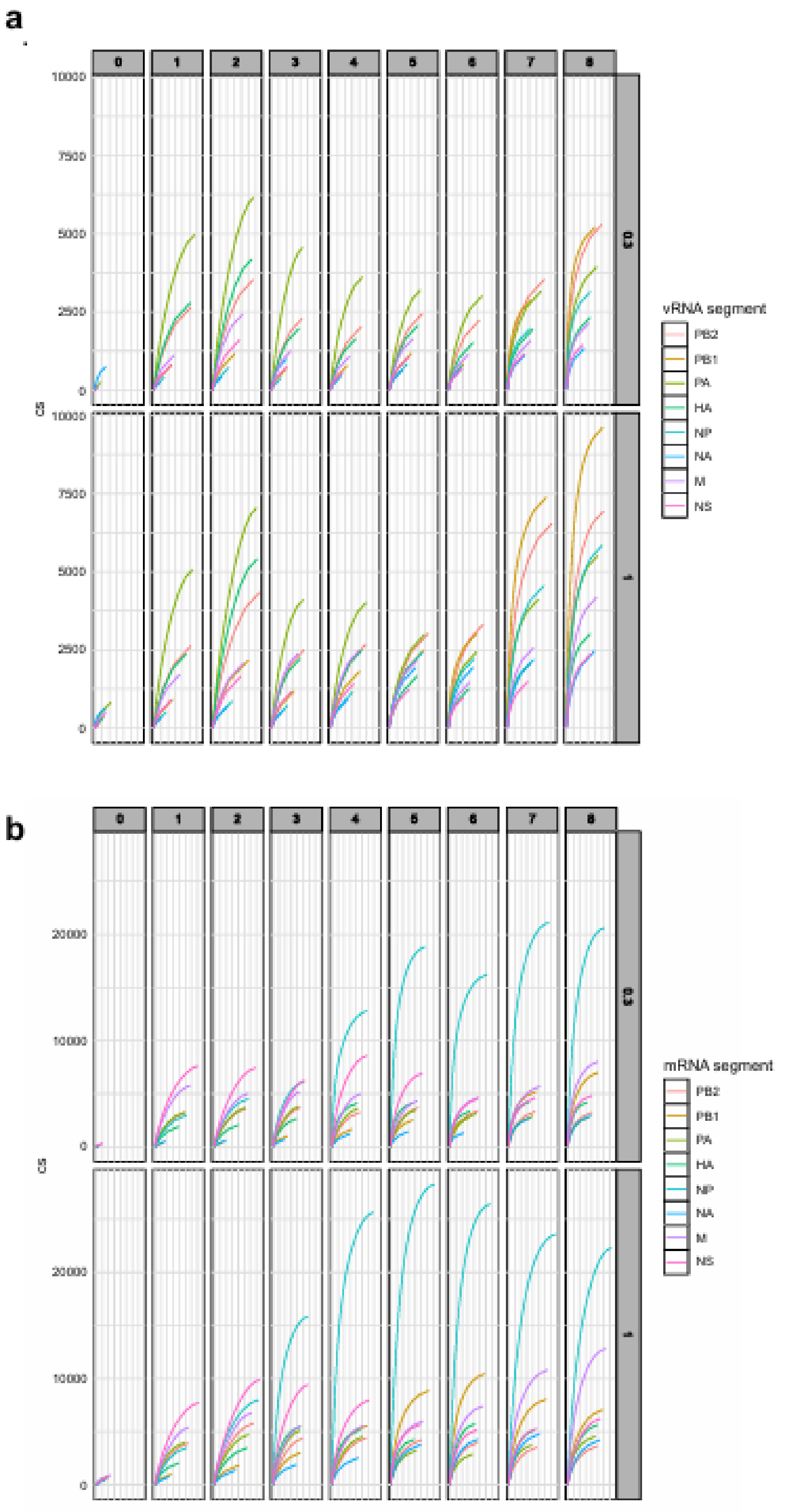

**Supplementary Figure 10. Single-cell heterogeneity in Influenza A virus RNA levels** (a, b) Distribution of copy numbers (cs) for each viral RNA (vRNA) and viral mRNA molecule in individual cells at 0-8 h post-infection for 0.3 and 1 MOI infected cells.

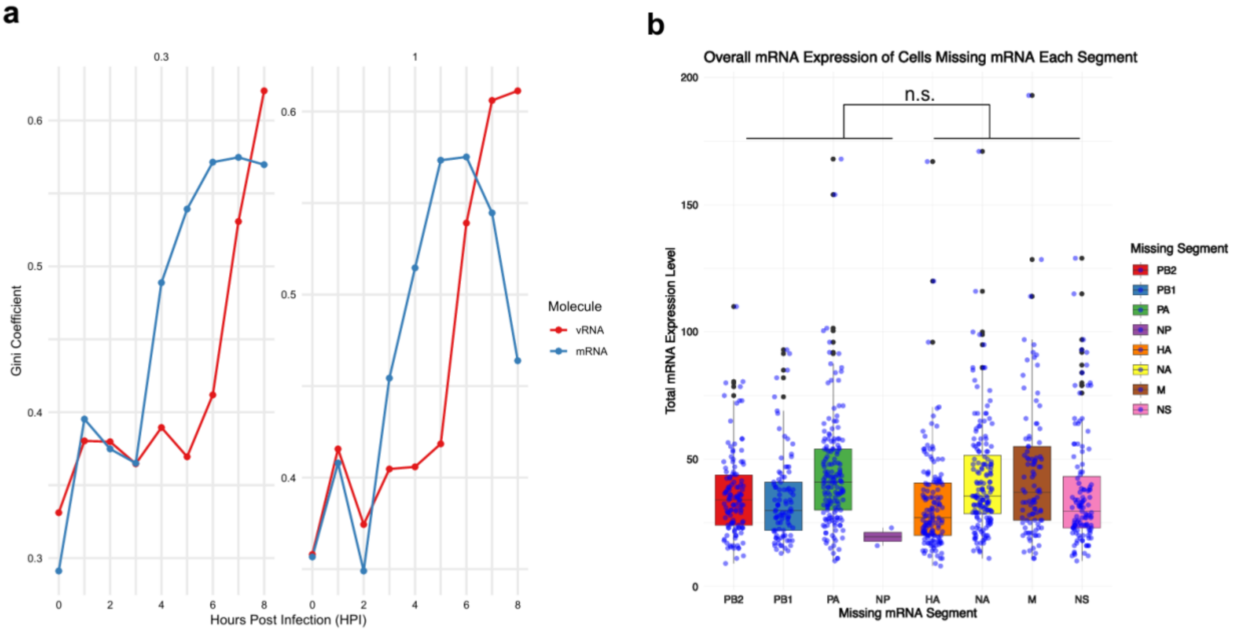


**Supplementary Figure 11. Quantification of Influenza A virus RNA distribution inequality.** (a) Gini coefficients for vRNA and mRNA distributions over time, quantifying the inequality in RNA copy numbers across the infected cell population for both 0.3 and 1 MOI infected cells. (b) Abundance of total mRNA in cells at hpi>5 missing exactly 1 mRNA segment. x-axis shows the missing mRNA segment, and the y-axis the total sum of mRNA in cells.

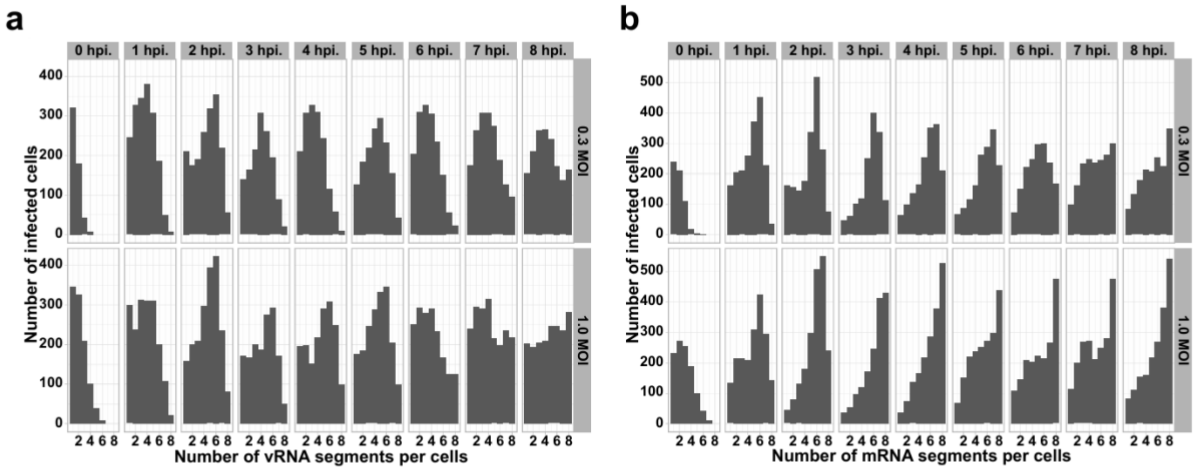


**Supplementary Figure 12. Analysis of complete and incomplete infection at single-cell level.** (a) Abundance of infected cells detected with one vRNA to complete 8 vRNA segments in infected cells, The x axis represents the observed number of segments (from just one segment to complete eight segments) and y axis represents the number of infected cells. (b)

Abundance of infected cells detected with one viral mRNA to complete 8 viral mRNA segments in infected cells, The x axis represents the observed number of segments and y axis represents the number of infected cells.

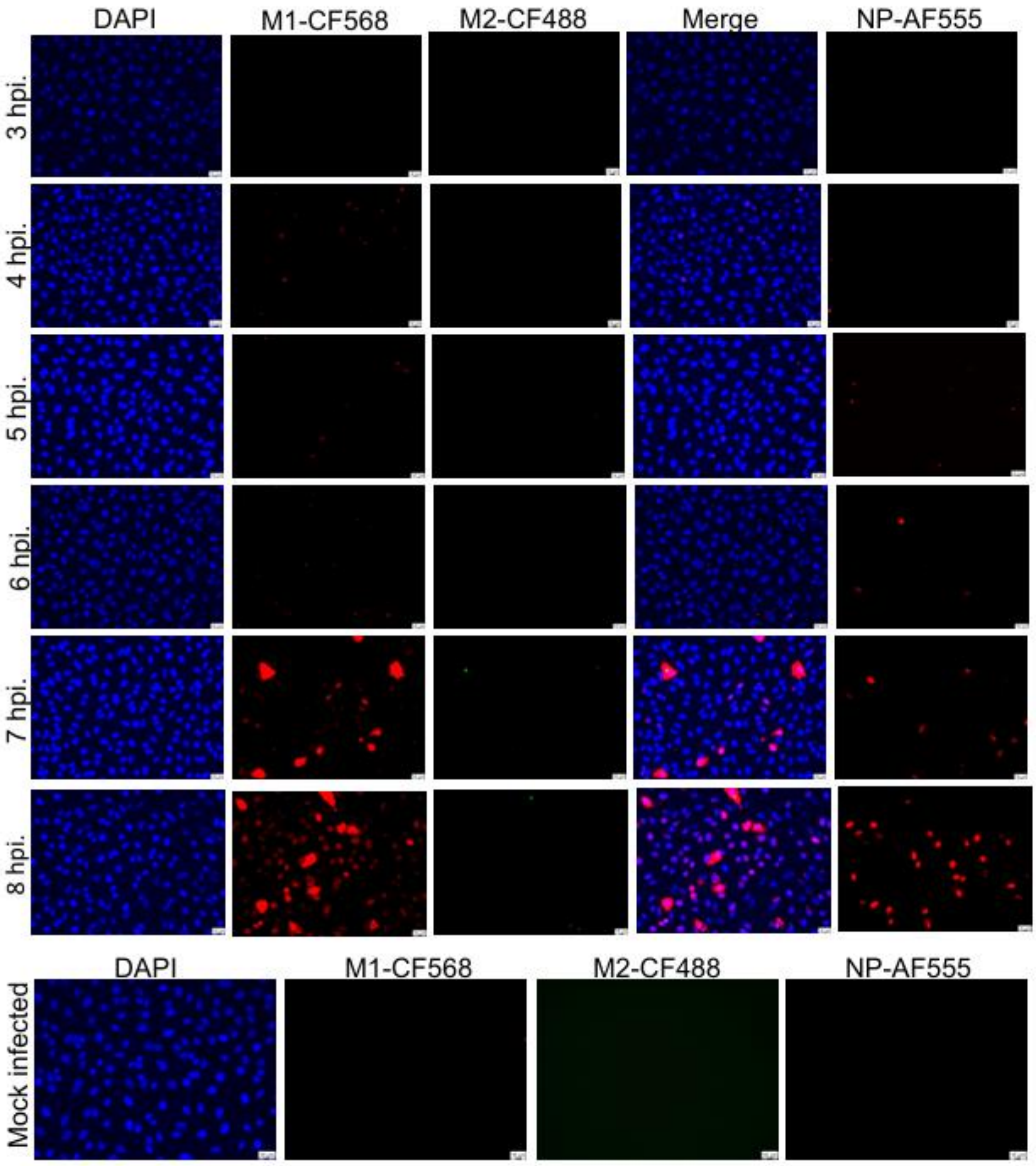


**Supplementary Figure 13. Immunofluorescence Imaging of MDCK Cells infected with PR8 Virus.** MDCK cells were infected with PR8 virus at a MOI of 0.3. After the indicated time points, cells were stained for viral proteins: M1 (red), M2 (green), and NP (red). Images are shown in individual imaging channels, including DAPI (blue) for the nucleus, along with separate channels for M1, M2, and NP. A merged image displays the nucleus (DAPI) along with M1 and M2 signals to visualize their localization.

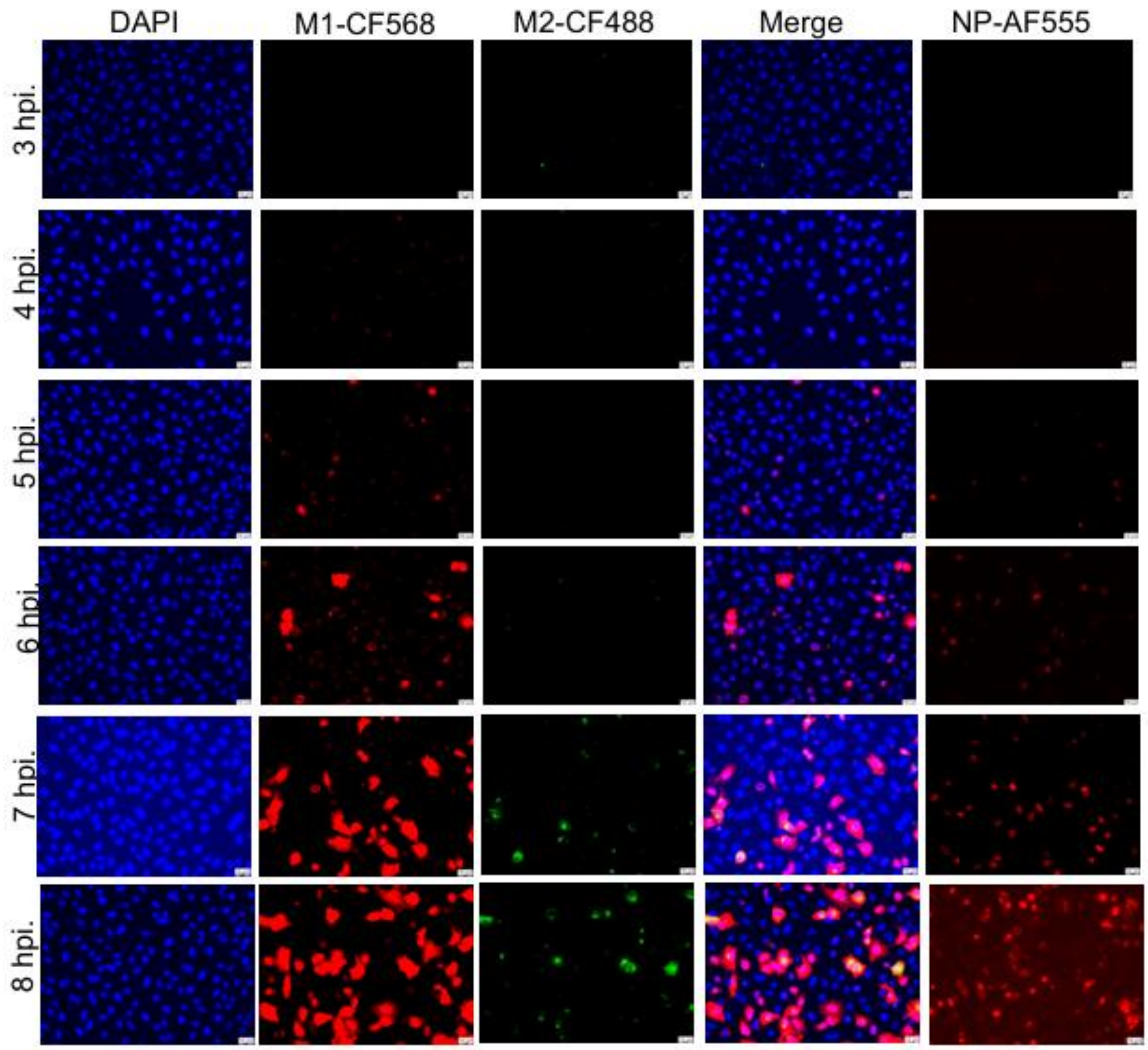


**Supplementary Figure 14. Immunofluorescence Imaging of MDCK Cells infected with PR8 Virus.** MDCK cells were infected with PR8 virus at a MOI of 1.0. After the indicated time points, cells were stained for viral proteins: M1 (red), M2 (green), and NP (red). Images are shown in individual imaging channels, including DAPI (blue) for the nucleus, along with separate channels for M1, M2, and NP. A merged image displays the nucleus (DAPI) along with M1 and M2 signals to visualize their localization.

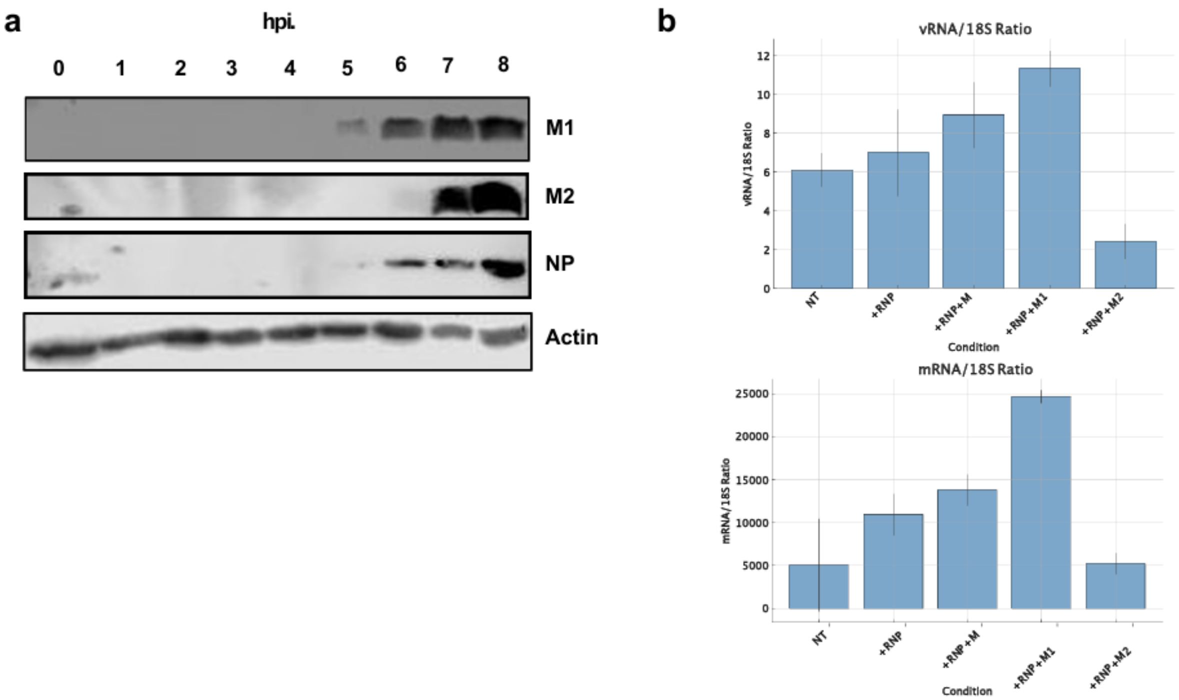


**Supplementary Figure 15. Time-dependent Expression of M1 and M2 in PR8-infected MDCK Cells.** (a) MDCK cells were infected with PR8 at an MOI of 5. At the indicated time points, the cells were harvested, and Western blot (WB) analysis was performed for M1, M2, NP protein. As a positive control actin protein was stained. M1 began to appear at 5 hpi, whereas M2 was first detected at 7 hpi. (b) Effect of RNP+M pre-expression on vRNA and mRNA levels in cells. RT-qPCR quantification of vRNA and vmRNA in cells pre-expressing RNP alone or in combination with M, M1, or M2.
